# Overexpression of the microtubule-binding protein CLIP-170 induces a +TIP network superstructure consistent with a biomolecular condensate

**DOI:** 10.1101/2021.01.01.424687

**Authors:** Yueh-Fu O. Wu, Annamarie T. Bryant, Nora T. Nelson, Alexander G. Madey, Gail F. Fernandes, Holly V. Goodson

**Author notes:** Corresponding author (HVG).

## Abstract

Proper regulation of microtubule (MT) dynamics is critical for cellular processes including cell division and intracellular transport. Plus-end tracking proteins (+TIPs) dynamically track growing MTs and play a key role in MT regulation. +TIPs participate in a complex web of intra- and inter-molecular interactions known as the +TIP network. Hypotheses addressing the purpose of +TIP:+TIP interactions include relieving +TIP autoinhibition and localizing MT regulators to growing MT ends. In addition, we have proposed that the web of +TIP:+TIP interactions has a physical purpose, creating a superstructure that constrains the structural fluctuations of the fragile MT tip and thus acts as a polymerization chaperone. Many animal +TIP network proteins are multivalent and have intrinsically disordered regions, features commonly found in biomolecular condensates. This observation suggests that the +TIP network might under some conditions form a biomolecular condensate. Previous studies have shown that overexpression of the +TIP CLIP-170 induces large “patch” structures containing CLIP-170 and other +TIPs. To test the hypothesis that these patches might be biomolecular condensates, we used video microscopy, immunofluorescence staining, and Fluorescence Recovery After Photobleaching (FRAP). Our data show that the CLIP-170-induced patches have hallmarks indicative of a biomolecular condensate, one that contains +TIP proteins and excludes other known condensate markers. Moreover, bioinformatic studies demonstrate that the presence of intrinsically disordered regions is conserved in key +TIPs, implying that these regions are functionally significant. Together, these results indicate that the CLIP-170 induced patches in cells are phase-separated liquid condensates and raise the possibility that the endogenous +TIP network might form a liquid droplet at MT ends or other +TIP locations.

## Introduction

Microtubules (MTs) compose one of the three major filament networks of the eukaryotic cytoskeleton, and they are required for basic cellular functions such as cell polarity, cell division and intracellular transport. Dysfunction of the MT cytoskeleton can lead to serious neurodegenerative diseases including tauopathies and Parkinson’s disease (1).

Microtubules display a surprising behavior known as dynamic instability, which describes the approximately random alteration between phases of slow growth (polymerization) and rapid shrinkage (depolymerization). This behavior is regulated by MT binding proteins and is central to MT function because it enables MTs to explore space to respond rapidly to internal and external signals and find organelles to be transported (reviewed by (2)).

The most conserved MT binding proteins (and by implication the most important) are a set of mutually interacting proteins that dynamically track growing MT ends and are collectively known as microtubule plus-end tracking proteins (+TIPs). +TIPs form an interaction network created by many weak, multivalent links (both intra- and inter-molecular) between MTs and +TIPs (2, 3). While many +TIPs and their MT regulatory roles have been identified, it is not yet fully understood why so many +TIPs bind to other +TIPs. One favored explanation is that the interactions of the +TIP network create regulatory pathways by relieving the autoinhibition feature present in many +TIP proteins (Fig 1A). Another explanation is that +TIP:+TIP interactions serve to localize and deliver proteins in a spatiotemporal manner (e.g. localizing +TIPs to the MT ends, and facilitating the surfing of proteins to cell edge) (reviewed in (3)).

**Fig 1.**
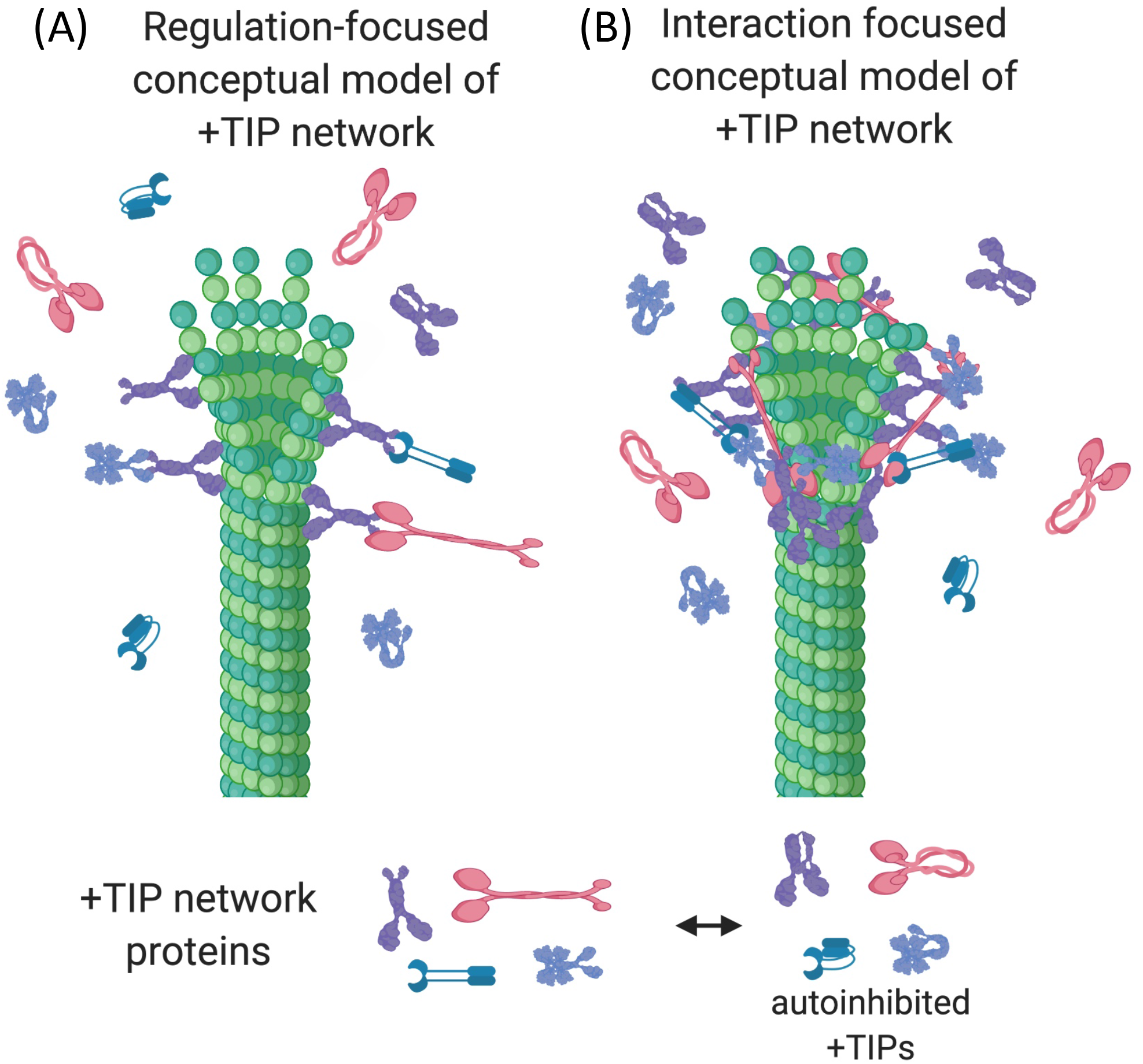
Conceptual models of the +TIP network. (A) In the regulation-focused model of the +TIP network, interactions between +TIPs function primarily to activate autoinhibited +TIP proteins and/or enable +TIPs to localize to the plus end. (B) In a more physical model of the +TIP network, interactions between +TIP proteins form a cross-linked, dynamic scaffold that supports and stabilizes the fluctuating MT tips by promoting lateral binding of protofilaments (4). This figure was created with BioRender (5).

While these explanations for the existence of the +TIP network are logical, some aspects of +TIP behavior seem inconsistent with either idea, suggesting this set of explanations is not complete. For example, deletion of evolutionarily conserved +TIPs often has surprisingly little effect on MT dynamics and/or cell survival (S1 Table), which is especially surprising if the localization and/or activity of some conserved +TIPs depend on others. In addition, while individual +TIPs can function independently in regulating MT dynamics, evidence from several labs also suggest that groups of +TIPs work synergistically to promote MT polymerization (e.g., (4, 6, 7)). Such synergy seems inconsistent with linear regulatory and/or localization pathways, which would predict epistasis or at most additive effects. These observations led us to suggest in earlier work that the interactions of the +TIP network have an important additional function, which is to create a superstructure that promotes MT assembly by constraining the structural fluctuations of the MT tip (Fig 1B) (4).

The work in this paper focuses on the +TIP CLIP-170, which was the first characterized +TIP (8). Previous studies have found that CLIP-170 promotes interactions between organelles and MTs (8, 9)(10) and regulates MT dynamics (11). CLIP-170 has also been implicated in actin-MT crosstalk (e.g., (12–14)). Similar to other +TIPs, CLIP-170 has multiple conserved binding sites that mediate interactions with MTs and/or other +TIPs (15, 16). Structurally, CLIP-170 contains three major domains. The N-terminal head domain is composed of two CAP-Gly subdomains and three serine-rich regions, all of which contribute to MT binding (15). The coiled-coil region mediates CLIP-170 dimerization (17) and contains a FEED domain, which forms a complex with formin and actin to promote actin polymerization (12). The C-terminal domain contains two zinc-knuckle motifs (9, 17, 18): The first of these mediates CLIP-170:CLIP-170 interactions involved in autoinhibition, while the second mediates an interaction with the dynein-associated dynactin complex (19, 20).

Overexpression of CLIP-170 causes formation of so-called “patches” (9, 19). The physical identity of these patches was puzzling to the researchers performing early work on CLIP-170, who initially assumed that the patches were membranous organelles of some kind but were not able to colocalize membrane markers to them. These early efforts showed that CLIP-170 could recruit other proteins more recently recognized as +TIPs (specifically EB1 and dynactin complex) into the patches (19). These characteristics and others described below led us to speculate that the patches might correspond to biomolecular condensates. The goal of the work described in this paper is to investigate this possibility and consider its potential implications in terms of the form and function of the +TIP network under normal physiological conditions.

Biomolecular condensates (aka liquid droplets, coacervates, membraneless organelles) are dynamic structures that spontaneously form when interactions between groups of weakly-binding biopolymers (protein and sometimes RNA or DNA) cause them to phase-separate from surrounding cytoplasm or nucleoplasm (21–24). Cytosolic examples include P-bodies, which regulate mRNA turnover (25, 26), Wnt signalosomes, which help to transduce Wnt signaling (27–29), and the pericentriolar material/matrix, which regulates MT nucleation at the centrosome (30, 31). In the nucleus, Cajal bodies regulate RNA-related metabolism (32), and PML bodies regulate nuclear functions (33). While these biomolecular condensates are associated with functions that promote cell survival, not all are positive: condensates of the microtubule-associated protein Tau corelate with the progression of Alzheimer’s disease (34–36).

The process in which proteins/nucleotides self-assemble into a condensate is called liquid-liquid phase separation (LLPS). This liquid-liquid demixing process becomes observable when the proteins/nucleotides reach a critical concentration (21–24). Biomolecules identified as forming condensates generally contain intrinsically disordered regions (IDRs) and multivalent binding regions that mediate weak interactions. These sequence features generate the driving forces for condensate formation (21–24). Physical factors such as temperature and pH can affect the condensation or dissolution of LLPS as well (21–24).

While many condensates are now well-recognized (21, 24), there are relatively few examples of mechanistically defined functions for condensate formation. Proposed functions include concentrating reactive molecules locally to promote cellular reactions (e.g., promoting actin nucleation (37)), serving as a storage depot (e.g., P-granules localized at the posterior of the embryo to produce germ cells (25)), and providing mechanical forces to generate local viscoelasticity, which can be important to resist the deformation of cellular structures (e.g. condensate formation at the centrosome (38)).

Even though the functions and localizations of these condensates are varied, some attributes are shared between them. Common hallmarks that are thought to identify a biomolecular condensate include a) elastic deformability, b) the ability to undergo fusion and fission, c) selectivity (i.e., inclusion of certain biomolecules and the exclusion of others), and d) rapid exchange of proteins between the droplets and the cytoplasm (21–23, 39).

Here, we use live-cell imaging, photobleaching, and immunofluorescence to show that the “patches” previously observed as being induced upon CLIP-170 overexpression have the four hallmarks of liquid-liquid condensates, leading us to conclude that the patches are biomolecular condensates. We also evaluated the sequence properties of CLIP-170 and other +TIPs for the presence of sequence characteristics associated with condensate formation. We show that many +TIPs both are multivalent and contain IDRs, and that for key +TIPs these sequence features are conserved across large spans of evolution, suggesting that they are functionally significant. These data are consistent with our previous proposal that the +TIP network forms a physical superstructure and lead us to speculate that this superstructure is (in at least some circumstances or locations) a biomolecular condensate that coats the MT tip to support and promote MT assembly.

## Materials and Methods

### Cell lines and cell culture

NIH3T3 cells (a gift of Dr. Reginal Hill) were grown in Dulbecco’s Modified Eagle Medium (DMEM, Sigma) plus 2% glutamine (BioWhittaker) and 10% bovine calf serum (VWR). Cos-7 cells (a gift of Dr. Kevin Vaughan) were grown in DMEM plus 1% glutamine and 10% fetal bovine serum (Sigma). Cells were incubated under standard cell culture conditions (37°C and 5% CO_2_).

### DNA constructs

CLIP-170 constructs utilize the version of CLIP-170 used in original Kreis lab publications (8), now described in accession AAA35693. CLIP-170 plasmids with expressed transgenes under control of the CMV promoter have been described previously: WT CLIP-170 in pSG5 vector (pSG5-myc-CLIP-170) (40), and the N-terminal EGFP-conjugated WT CLIP-170 in pCB6 vector with EGFP (pCG6-CB6-CLIP-170, “GFP-CLIP-170” here after) (41).

### Antibodies & lipid dyes

The CLIP-170 antibodies used in this study were either a rabbit polyclonal raised against *Xenopus* CLIP-170 (this manuscript) or the mixture of mouse monoclonal antibodies 4D3/2D6 (8, 42), with the exception of the GalT colocalization experiment, which utilized the rabbit polyclonal ɑ55 raised against human CLIP-170 (8). p150 was recognized by rabbit polyclonal anti-p150^Glued^, which was a gift from K. Vaughan and R. Vallee. EEA1 was recognized by rabbit polyclonal anti-EEA1 (43). The monoclonal antibody against GalT was a gift to Dr. Thomas Kris from Dr. T. Suganuma (44). Other primary antibodies were obtained from commercial sources: hnRNPA1 - mouse hnRNPA1 monoclonal antibody 4B10 (Invitrogen, MA5-24774), YB1 - rabbit YB1 polyclonal antibody D299 (Cell signaling, 4202S), Staufen1 - rabbit polyclonal antibody (Bios antibodies, bs-9877R), CLASP2 - rabbit polyclonal antibody (Proteintech, 12942-1-AP), mDia2 - rabbit DIAPH3 polyclonal antibody (Proteintech, 14342-1-AP), EB1 - mouse monoclonal antibody (Transduction Laboratories, now BD-Transduction, 610534), and LIS-1 - mouse monoclonal antibody H-7 (Santa Cruz, sc-374586). All labeled secondary antibodies were obtained from Invitrogen. Generic RNA was stained with SYTO™ RNASelect™ Green Fluorescent Cell Stain (Thermo Fisher, S32703). Rhodamine DHPE (Thermo Fisher, L-1392), Dil (Invitrogen), and RhPE (Molecular Probes) were used as generic membrane labels. LysoTracker (Molecular Probes, DND-99) and MitoTracker Red CMXRos (Cell Signaling) were used to stain lysosome and mitochondria, respectively. For all membrane staining experiments, cells were stained according to manufacturer protocols, and then moved into imaging medium for imaging.

### Transient transfections

For fixed cell images, cells were grown on 10 mm^2^ glass coverslips (Knittel Glaser). For live cell imaging, cells were grown on 35 mm dishes with 10 mm glass bottom coverslips (P35G-1.5-10-C, MatTek) or 8-well glass-bottom chambers (Eppendorf). Cells were transiently transfected by Lipofectamine 3000 (Invitrogen, L3000008) with 5 μg of indicated construct DNA per reaction when cells reached 70-80% confluence. For consistency, cells were incubated for 24 hr post-transfection before performing live imaging, FRAP, and immunofluorescence assays.

### Immunofluorescence

As indicated in the figure legends, cells were fixed by methanol or 3% paraformaldehyde (PFA) before staining with the antibodies described above. For the methanol fixation, cells were fixed in −20°C methanol for 4 min and washed three times with PBS before treatment with primary antibodies. For the PFA fixation, cells were fixed in 3% PFA for 20 min, washed once with PBS, quenched with a wash of 30 mM ammonium chloride in PBS, and washed once again in PBS. Cells were then permeabilized with 0.1% Triton-X-100 for 4 min and washed with PBS again before treating with primary antibodies. Each washing step was 5 min. Cells were incubated with the primary antibody for 20 min, washed three times with PBS, incubated with the indicated fluorophore-conjugated secondary antibody for 20 min, and washed three times with PBS. Cells were mounted on slides in Mowiol 4-88 mounting medium (Sigma 475904-M). Unless otherwise indicated, images were acquired with a 100x objective (1.4 N.A.) and a 1.5× optivar on a TE2000 inverted microscope (Nikon) with a Hamamatsu CMOS camera controlled by NIS-Elements BR 413.04 64-bit software (Nikon). Alternatively, microscopy was performed on a DeltaVision Deconvolution Microscope (GE Healthcare) with a 60x 1.42 N.A. objective connected to a Photometrics Coolsnap hq2 monochrome camera controlled by Resolve3D – SoftWoRx 7.0.0 software (GE Healthcare). Images were processed using Fiji (National Institutes of Health, https://imagej.net/Fiji) (45). For the EEA1 and GalT immunofluorescence experiments in Supplementary Information, cells were imaged using a 40x 1.4NA PlanApo objective on Zeiss inverted microscope (TV135); images were recorded with a cooled CCD camera (Photometrics CH250, Tuscon AZ) driven by IPLab-Spectrum software (Scanalytics, Fairfax VA), and processed by Photoshop (Adobe). Because CLIP-170 patches can be very bright, negative controls with one primary and two secondary antibodies were routinely performed to assess levels of signal created by bleed through and/or antibody cross-reactivity.

### Live imaging and Fluorescence Recovery After Photobleaching (FRAP)

Live imaging experiments were performed either with an A1R-MP Laser Scanning Confocal Microscope or with DeltaVision as indicated. In both systems, cells were incubated at 37°C with 0.5% CO_2_ within an environmental chamber. DMEM culture media lacking phenol red (Sigma, 51449C) plus 2% glutamine and 10% bovine calf serum (VWR) was used during live imaging processes.

For the FRAP measurements, FRAP experiments were performed on the A1R-MP Laser Scanning Confocal Microscope (Nikon) using a 100x 1.49NA oil immersion objective, which was driven by Nikon NIS-Elements software and was equipped with PMT detectors. Condensates were bleached with a solid state laser (Coherent) with a 488 nm, 100% laser intensity for 200 ms. Time-lapse images were acquired before and after photobleaching at two frames per sec for 9 loops and 82 loops, respectively. We set the bleach regions as a 1 μm diameter circle, though the region that was actually affected by the bleaching was ~1.5-2 μm in diameter.

Images were processed using ImageJ to measure the intensity of the regions of interest (FRAP region, background region, and the reference region) before and after photobleaching. The selection of regions of interest and FRAP analysis were performed following the guide of the protocol of the Aravin lab and EMBL (46). Briefly, we measured the intensity of three different regions to assess photobleaching: (1) BL: the region that was bleached by the laser, (2) BG: a region outside of the cell, used to measure the level of background signal, and (3) REF: a region inside of the cell but outside the bleached region, used to assess the decay of signal from imaging-associated photobleaching. After determining the intensities of the three regions over time, we subtracted the background signal (BG) from the BL and REF regions and normalized the fluorescence decay by dividing BL by REF to get the normalized intensity over time I(t): 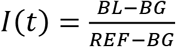. Next, to calculate the fraction of fluorescence recovery after photobleaching, we divided the intensity after bleaching by the mean intensity before bleaching to get: 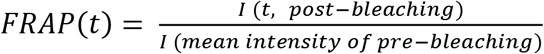. Plotting FRAP(t) over time gave an intensity curve such as that shown in (Fig 5B). Finally, we used MATLAB(R2019a) to fit the intensity curve with the equation *FRAP*(*t*) = *a* * (1 − *e*^−*bt*^) + *c*, where a is the slowly recovering fraction; b is recovery rate constant; c is the rapidly diffusing fraction (46–49). The fraction of mobile protein = a+c, and the fraction of immobile protein = 1-(a+c). The halftime of recovery (t_1/2_) = ln2/b.

### Bioinformatics

#### Identification of +TIPs in other organisms

A range of nine model organisms (from plants to humans) was chosen to assess the conservation of the presence of intrinsically disordered regions (IDRs) in a set of well-recognized +TIP network proteins. The selected organisms are: *Homo sapiens* (taxid:9606), *Mus musculus* (taxid:10090), *Xenopus laevis* (taxid:8355), *Danio rerio* (taxid:7955), *Drosophila melanogaster* (taxid:7227), *Saccharomyces cerevisiae* (taxid:4932), *Schizosaccharomyces pombe* (taxid:4896), *Dictyostelium discoideum* AX4 (taxid:352472), and *Arabidopsis thaliana* (taxid:3702).

To identify core +TIPs (or other relevant proteins) in these organisms, we performed a BLASTp (or when necessary, a PSI-BLAST) search in the NCBI RefSeq database using the following representative human sequences as probes: CLIP-170 (AAA35693.1), EB1 (NP_0364571), MAP215 (NP_0010089381), CLASP2 (NP_0013525571), CLASP1 (NP_0560971), p150 (NP_0040732), LIS-1 (NP_0004211), APC (NP_0013418251), and mDia2 (NP_0010359821). In many cases (e.g., EB1), identification of unambiguous homologs was straightforward from examination of the e-values and alignments; in cases where gene duplications had occurred, we chose the gene with the strongest (lowest) e-value for further analysis. In cases of alternative splicing, we chose the longest of the curated transcripts for further analysis, except for CLIP-170 and CLASP homologs, where we chose the form that best matched the domain structure of the human protein used as the probe. Note that CLIP-170 homologs were not identified in *Dictyostelium* or *Arabidopsis*.

#### Bioinformatic tools used

##### IDR prediction

We focused our efforts on three IDR predictors, which were used with default parameters unless otherwise indicated: 1) “ESpritz version 1.3” (50) (http://old.protein.bio.unipd.it/espritz/), used with prediction type as X-Ray and decision threshold as 5% false positive rate (5% FPR; chosen because the default setting [“Best Sw” – a weighted score rewarding prediction of IDRs] tends to overestimate IDRs); 2) “Interpro” (EMBL, https://www.ebi.ac.uk/interpro/), which provides as part of its standard analysis disorder predictions from the MobiDB-lite database (51)); and 3) “IUPred2”, used with “short disordered” setting, as obtained from the IUPred2A server (https://iupred2a.elte.hu/) (52, 53). We chose these three out of the wide array of possible IDR predictors because they are widely-used (54), and in our preliminary analysis they rarely generated spurious reports of coiled-coil regions as IDRs, unlike some others, e.g., PONDR. In Fig 6, we present the results of all three methods for human CLIP-170; for other analyses, we simplified the display by showing only the results from the first two predictors because the results from Espritz and IUPred2 were similar. The likelihood of intrinsic disorder as a function of amino acid position as produced by ESpritz and IUPred2 were plotted by MATLAB(R2019a). For MobiDB-lite predictions, we manually listed all predicted IDRs as obtained from Interpro. Then, we plotted the list of IDRs with the MATLAB(R2019a) “patch” function.

##### Other analyses

Sets of orthologous protein sequences were aligned by ClustalW (55) or alternatively PASTA (56). For most of the domain structure predictions, we used a hmmscan Pfam search with e-value 0.1 at HMMER (https://www.ebi.ac.uk/Tools/hmmer/search/phmmer) (57) (S5-S8 Figs). The only exception was CLIP-170 and its homologs. To map the domain structure of CLIP-170 homologs, we extracted from the NCBI protein information page the location of CAP-Gly domains, serine-rich regions, and zinc-knuckle motifs. Then, we aligned the human CLIP-170 FEED domain (VEEESITKGDLE) (12) to the representative sequences of other organisms to identify the FEED domain. Lastly, we combined all the domain information and plotted the protein sequence with “IBS Version 1.0” (http://ibs.biocuckoo.org/online.php). To predict coiled-coil regions, we used the “COILS” predictor (https://embnet.vital-it.ch/software/COILS_form.html) (58) with default settings.

## Results

The overall goals of this study were to test whether the so-called “patches” induced by CLIP-170 overexpression (9, 19) are biomolecular condensates and, if so, to then consider the implications of this observation for the structure and function of the +TIP network under normal physiological conditions. Although it would be ideal to examine the endogenous +TIP network directly, the “comets” at the ends of MT plus ends are both dynamic and small (near the diffraction limit of visible light), making it technically difficult to determine clearly whether they have condensate properties. Thus, although CLIP-170 patches are themselves overexpression artifacts, we hypothesized that they might reflect some properties of physiological +TIP networks and so used them as a tool to gain insight into these properties. As evidence that this approach can be fruitful, in previous work we successfully used this same logic to identify the +TIPs EB1 and dynactin complex as CLIP-170-interacting proteins and showed that the interactions were mediated by different regions of the CLIP-170 protein (19).

As noted above, four properties common to biomolecular condensates are 1) elastic deformability, 2) the ability to undergo fusion and fission, 3) selectivity: droplets include certain members but exclude others, and 4) rapid exchange of proteins between the droplets and the cytoplasm (21–23, 39). Thus, we examined the CLIP-170-induced patches for each of these characteristics in turn. In addition, proteins included in droplets generally have the following structural characteristics that are important for droplet formation: they have multiple sites for binding other droplet members (i.e., they are multivalent), and they contain intrinsically disordered regions (IDRs). Accordingly, we also examined CLIP-170 and other +TIPs for these characteristics.

### The so-called “patches” induced upon CLIP-170 overexpression do not colocalize with membranes

As shown previously, when CLIP-170 is expressed at low levels, it labels MT plus ends, but at higher levels causes formation of patches and microtubule bundles (Fig 2B, (19)). This phenomenon is seen in a range of cell types, including HeLa (9, 19), Cos-7 ((19), Fig 2B), and NIH3T3 (Fig 2B). To test whether the patches are membraneless, we first examined the patches for colocalization with various specific membrane markers, including MitoTracker, LysoTracker, ER tracker, GalT (labels Golgi), and EEA1 (labels endosomes). None of these, with the possible exception of EEA1, showed any significant colocalization with the CLIP-170 patches (S1 Fig).

**Fig 2.**
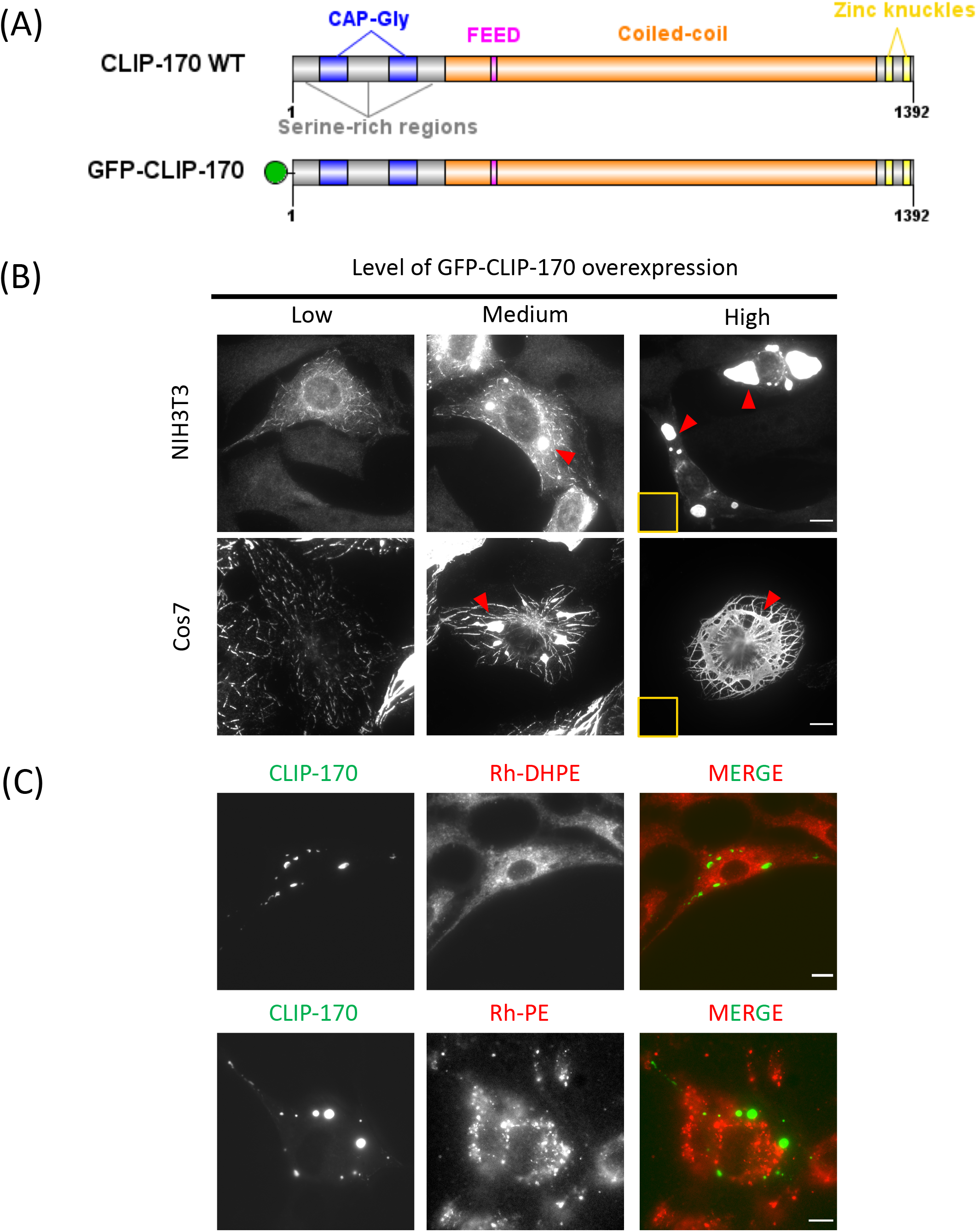
CLIP-170 patches do not colocalize with membrane markers. (A) Domain structures of the CLIP-170 constructs used in this study. (B) NIH3T3 or Cos-7 cells (as indicated) were transfected for 24 hr to overexpress GFP-CLIP-170, fixed with PFA, and observed by widefield fluorescence microscopy. Arrows point out examples of CLIP-170 patches as observed in cells at varied expression levels as indicated. The contrast of each image in a given row was adjusted to the same levels, except for high-level expression, where the main image was adjusted to allow the visualization of large and bright patch structures, and an inset provides a version normalized to match the rest of the row. (C) As an example of the failure of membrane markers to colocalize with patches, NIH3T3 cells were transfected with GFP-CLIP-170 for 24 hours. Rhodamine-DHPE or Rhodamine-PE was used as a generic membrane label to stain lipid membranes. Live cells were observed by widefield fluorescence microscopy. See S1 Fig for colocalizations with additional membrane markers. Scale bar: 10 μm

One problem with using specific makers to test for membrane presence is that cells contain many different membrane-bound compartments, and so negative results are not definitive: we might simply have failed to test for the appropriate compartment. Therefore, we tested for the presence of membranes more generally by looking for colocalization with the nonspecific membrane dyes Rhodamine-DHPE and Rhodamine-PE. We observed no obvious colocalization with either dye (Fig 2C). These results are consistent with the hypothesis that the CLIP-170-induced patches are membraneless biomolecular condensates. Therefore, we proceeded to test for the presence of other characteristics of biomolecular condensates.

### CLIP-170 patches can elastically deform and undergo fusion and fission

As discussed above, another characteristic of biomolecular condensates is that they can undergo elastic deformations, fission, and fusion. To test whether CLIP-170 patches exhibit these activities, we overexpressed GFP-CLIP-170 in NIH3T3 cells and used live cell fluorescence microscopy to monitor the GFP-CLIP-170 behavior over time. As shown in Fig 3 and Movie 1, CLIP-170 patches do indeed display fusion events (red box) and fission events (magenta arrow). In addition, it is common to see CLIP-170-labeled +TIP comets travel through and deform the CLIP-170 patches (Fig 3 and Movie 1). This observation indicates that the CLIP-170 patches interact with the +TIP network and suggests that these two structures may share some properties with each other. Taken together, these results indicate that the CLIP-170 patches have the elastic deformability, fusion and fission properties expected of a biomolecular condensate.

**Fig 3.**
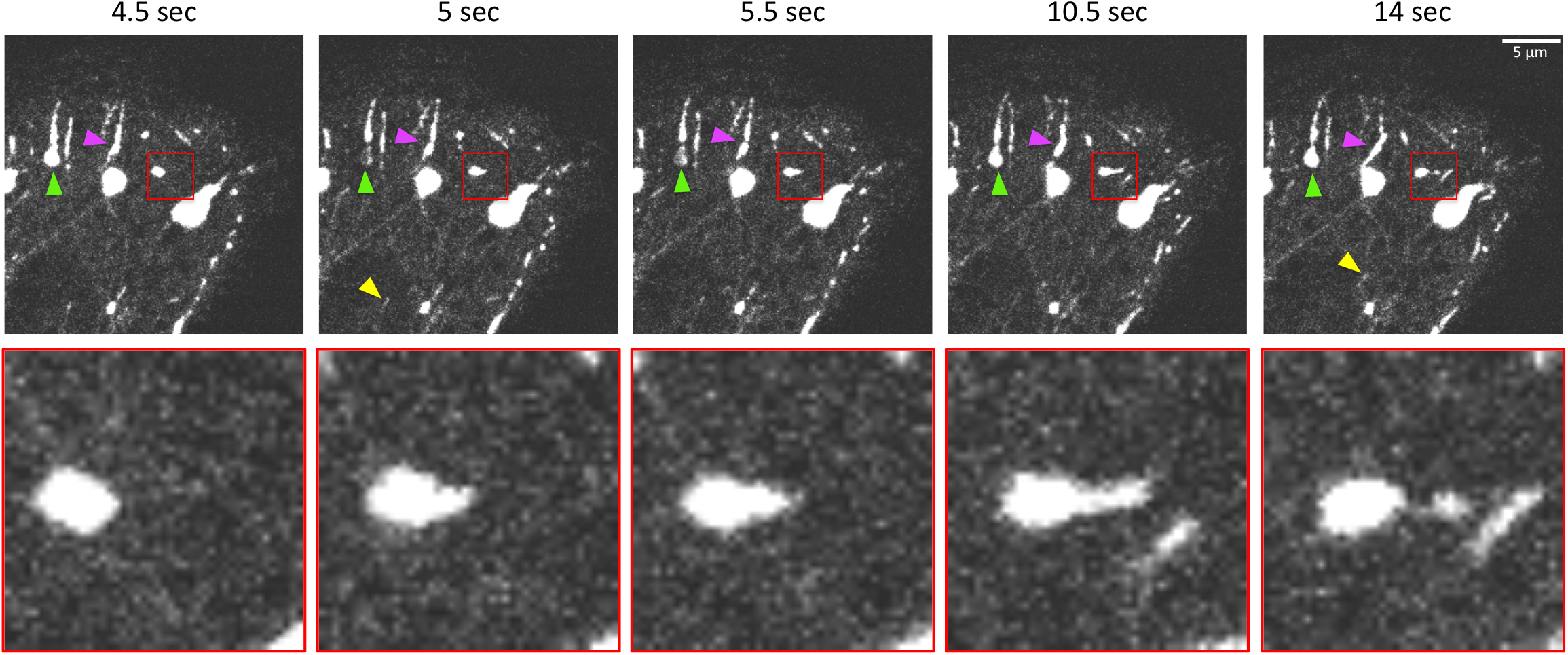
Dynamic behaviors of CLIP-170 patches *in vivo*. NIH3T3 cells were transfected to overexpress GFP-CLIP-170, and the behavior of GFP-CLIP-170 *in vivo* was recorded by confocal microscopy after 24-27 hr of transfection. Red box: An example of an apparent elastic deformation of a patch, following by fission (blow-up of this region is shown below). Arrows indicate examples of patch fusion (magenta), a photobleached site (green, relevant to discussions below), and a typical comet-shaped +TIP localization on a MT tip (yellow).

### CLIP-170 patches selectively recruit some cytoskeletal proteins but exclude members of other biomolecular condensates

The next biomolecular condensate property we tested was selectivity: do CLIP-170 patches selectively recruit some proteins and exclude others? To address this question, we used immunofluorescence staining to test various molecules for patch colocalization. The two groups of markers we tested were: (1) members of the +TIP network and (2) proteins included in established cytosolic liquid droplets (Table 1).

**Table 1.** List of molecules we have tested for co-localization with patches

| **Protein** | **Included/excluded** | **Protein marker of** |
| --- | --- | --- |
| **EB1** | Included | +TIPs |
| **LIS-1** | Included | +TIPs |
| **APC** | Excluded | +TIPs and Wnt signaling |
| **CLASP1** | NT | +TIPs |
| **CLASP2** | Excluded | +TIPs and Golgi |
| **XMAP215** | NT | +TIPs |
| **p150** | Included | +TIPs and Dynactin |
| **mDia2** | Included | Formin |
| **RNA** | Excluded | RNP granules |
| **YB1** | Excluded | Stress granules and P-bodies |
| **hnRNPA1** | Excluded | RNP granules |
| **Staufen1** | Excluded | RNP granules |

Experiments were conducted in NIH3T3 cells. NT indicates “Not Tested” due to lack of suitable antibodies.
See Fig 4 and S2 Fig for images of the colocalizations summarized above.

As expected from previous work (19), the +TIPs EB1 and p150 dynactin were observed to be recruited to the CLIP-170 patches (Fig 4, Table 1). In addition, we found that the patches recruit the +TIP LIS1, as well mDia2, an actin nucleator that binds EB1 (59) and so might be described as a peripheral part of the +TIP network (Fig 4, Table 1). However, antibodies against some other MT binding proteins including the +TIP CLASP2 did not colocalize with CLIP-170 patches (S2 Fig, Table 1).

**Fig 4.**
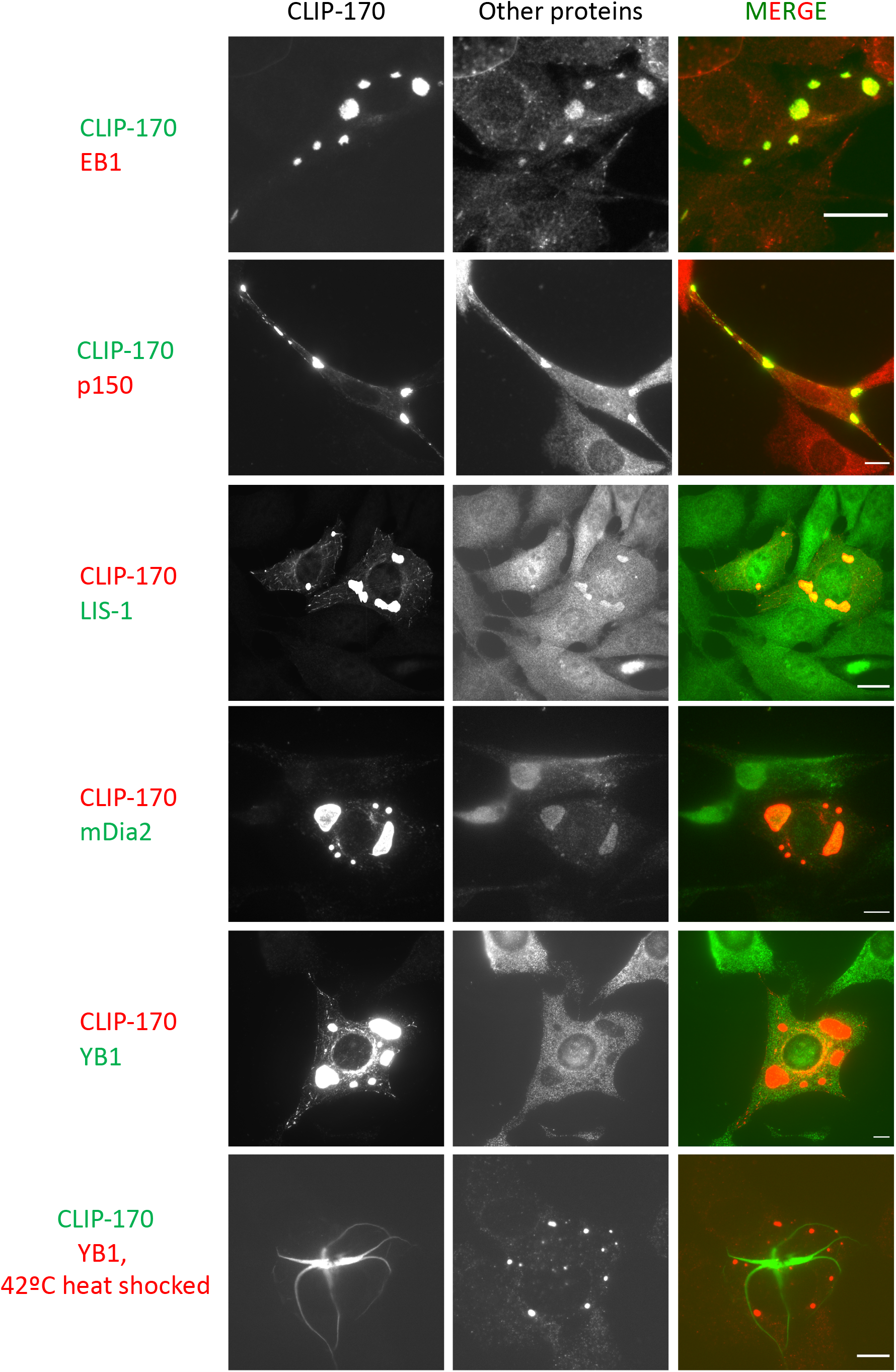
CLIP-170 patches have selective properties. NIH3T3 cells were transfected with full-length CLIP-170 for 24 hr before fixation and staining as indicated. For the EB1, p150, and YB1 labeling, cells were fixed with methanol; for LIS1 and mDia2, cells were fixed with PFA. The contrast of all images in a given row was set to the same levels. These data show that EB1, p150, LIS-1, and mDia2 were included in the CLIP-170 patches; YB1 is an example of a molecule that is excluded from the patches. See S2 Fig for images of colocalizations with other molecules, and see Table 1 for summarized list. Scale bar: 10 μm.

**Fig 5.**
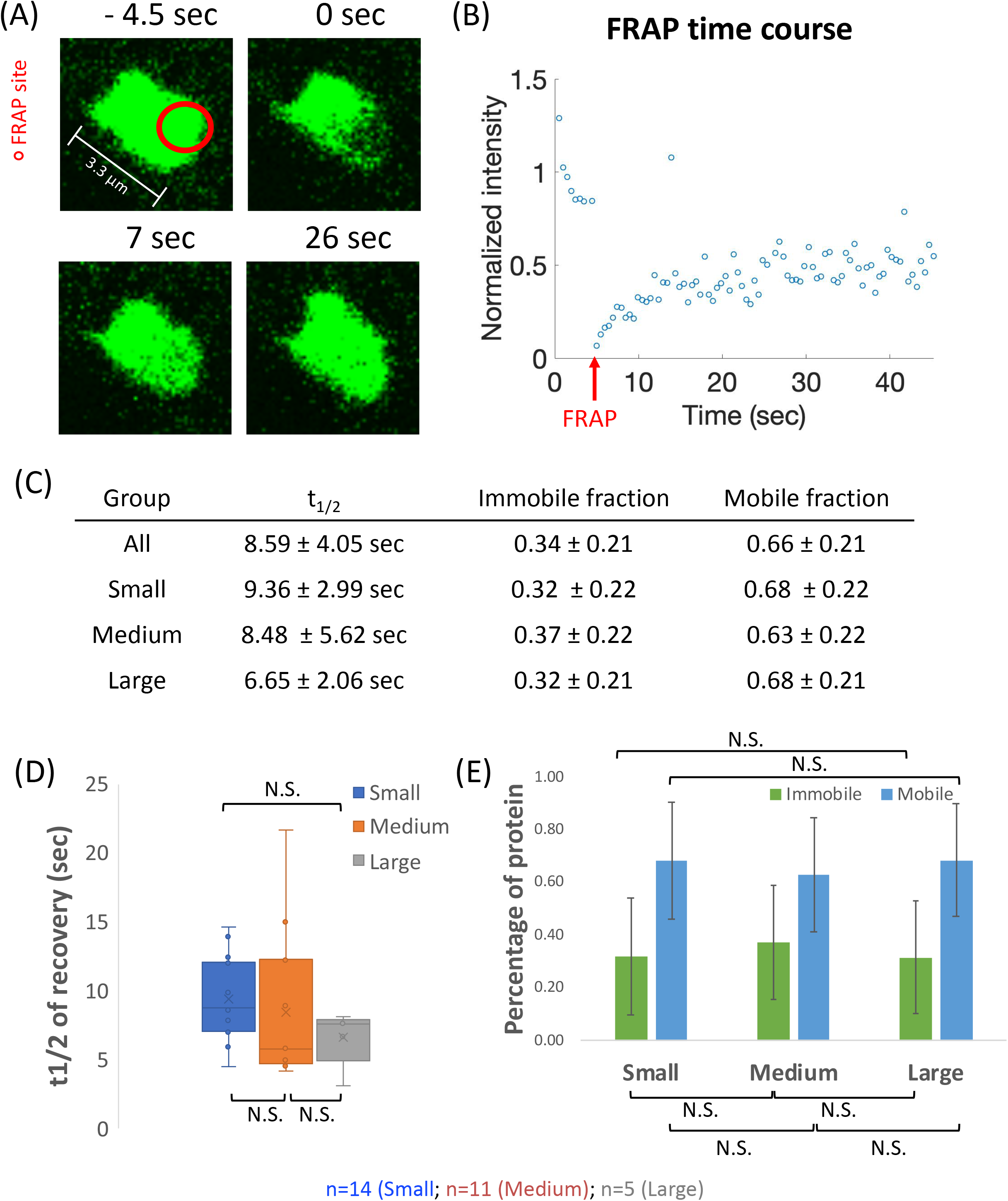
FRAP analysis of the dynamics of CLIP-170 patches. (A) Example of a partially photobleached CLIP-170 patch in an NIH3T3 cell. (B) The time course plot for the FRAP spot in (A) represents the fluorescence recovery of the bleaching site. Data were normalized against the average mean intensity before bleaching. (C, D, E) Quantification of the t_1/2_, mobile fraction, and immobile fraction. A total of 30 condensates were photobleached, and their recovery profiles were fitted to the exponential recovery equation (see Methods for details). “All” indicates the summation of data without regard to droplet size. In addition, we analyzed the droplets as separated into small, medium, and large groups (see S3 Fig for more information about how this separation was performed). (C) provides a numerical summary of the t_1/2_, mobile fraction, and immobile fraction. Panel (D) provides a box plot of the t_1/2_ of each size group, while panel (E) provides a bar graph of the fractions of mobile and immobile protein. n=14 (Small), 11 (Medium), and 5 (Large), for a total of 30 droplets. The error bars show the standard deviation; other aspects of the box plot are as generated by the default Excel Box and Whisker plot function. N.S. = Not Significant (there were no significant differences between any of these groups).

**Fig 6.**
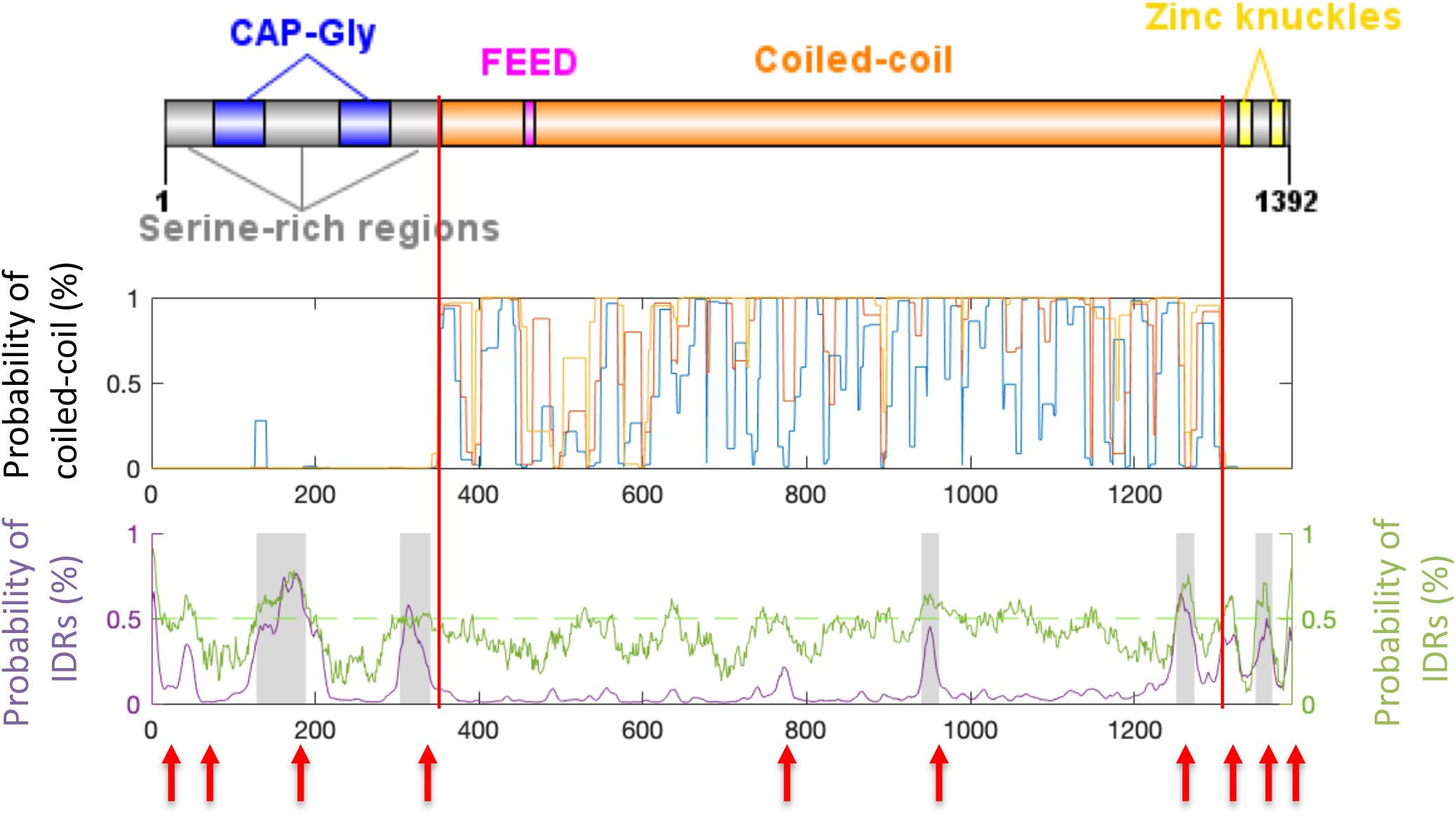
Analysis of coiled-coil domains and IDRs for human CLIP-170 (AAA35693.1). (Top) CLIP-170 domain structure summary. The N-terminal domain includes the 2 CAP-Gly domains and 3 serine-rich regions. The coiled-coil domain is labeled in orange, and its boundaries are indicated with two red lines. The C-terminal domain contains two zinc-knuckle motifs (yellow). (Middle) Analysis of coiled-coil propensity as predicted by COILS. The colored lines represent the probability that a region assumes a coiled-coil conformation, as assessed for different windows (7AA, blue; 14 AA, orange; 21 AA yellow), with 1 indicating 100% likelihood. (Bottom) IDR predictions (Y-axis) as function of amino acid position (X-axis), with 1 indicating 100% likelihood. The purple line represents the probability of IDRs as predicted by Espritz, with red arrows indicating the 10 regions predicted to be disordered. The grey areas indicate the IDRs as predicted by MobiDB-lite. The green line represents the probability of IDRs as predicted by IUPred2A, and the light green indicates the probability of 0.5.

To test for possible overlap between the CLIP-170 patches and other biomolecular condensates, we checked markers of RNP granules, P-bodies, and stress granules for colocalization to the patches. None were included in the CLIP-170 patches under our normal culture conditions (Figs 4, S2 Fig, and Table 1). Because stress granules are typically visible only under stress, we also tested colocalization with the stress granule proteins YB1 and Staufen1 after 42°C heat shock; the heat shock did induce the formation of structures containing these proteins, but they did not colocalize with the CLIP-170 patches (Figs 4, S2 Fig, and Table 1).

Taken together, these observations demonstrate that CLIP-170 patches selectively include a specific set of +TIP network proteins while excluding members of other biomolecular condensates, again consistent with the idea that the patches are biomolecular condensates.

### CLIP-170 in the patches exchanges rapidly within the cytoplasm

Another expected characteristic of a biomolecular condensate is the rapid molecular exchange of molecules between the condensate and the cytoplasm. A technique commonly used to assess such exchange is Fluorescence Recovery After Photobleaching (FRAP). We performed FRAP assays on the CLIP-170 patches and observed that the fluorescence intensity of the photobleached spots recovered quickly (half time of recovery (t_1/2_) = 8.59 ± 4.05 sec) and to a large degree (0.66 ± 0.21 mobile fraction) (see example in Fig 5). This rate and degree of recovery of CLIP-170 patches is consistent with those reported for other biomolecular condensates (60).

To further understand the kinetics of CLIP-170 patches, we categorized our FRAP data into three groups by patch diameter to determine whether the fluorescence recovery time and/or degree varied with size: small (0 – 2.4 μm), medium (2.4 – 4.4 μm), and large (≥ 4.4 μm). The grouping strategy is based on the size distribution after measuring the diameters of ~300 condensates (S3 Fig).

We observed that while the recovery of the medium patches was more variable than that of large and small patches, there was no significant difference between the groups in either the recovery rate or recovery extent (Figs 5C-E). These observations are notable for three reasons. First, since bleaching generally covered the full area of the small patches, the observation that small patches recover quickly confirms that the patches are not membrane bound because the interior of a membrane-bound compartment would not be expected to exchange material with the cytoplasm (exterior) on this time scale. Second, the observation that all three patch sizes recover with a similar rate indicates that the surface area: volume ratio is not rate limiting for material exchange with the cytoplasm. This in turn suggests that the patches are relatively porous. Third, the observation that the size of the condensates does not obviously correlate with the observed exchange dynamics is consistent with the idea that the CLIP-170 patches may reflect processes occurring in the small, endogenous +TIP network comets (~0.5 μm).

Taken together, our results thus far demonstrate that the CLIP-170 patches share key properties with known biomolecular condensates: they are membraneless, they undergo elastic deformations, fission, and fusion, they selectively include some molecules (e.g., +TIPs) while excluding others (e.g, stress granule proteins), and proteins exchange rapidly between patches and the cytoplasm. These observations lead us to conclude that the CLIP-170-induced patches are biomolecular condensates.

### Sequence information is consistent with the idea that CLIP-170 is a component of biomolecular condensates

If CLIP-170 is inducing biomolecular condensates, one would expect CLIP-170 to have sequence attributes consistent with this activity. As mentioned above, proteins found in condensates (i.e., proteins that have the potential to undergo phase-phase separation) typically are multivalent (i.e., have multiple binding sites for other biomolecules) and contain intrinsically disordered regions (IDRs). Indeed, it is already well-established in the literature that CLIP-170 has multivalent interactions with multiple other proteins: a dimer of CLIP-170 has as many as ten binding sites for tubulin (15), multiple binding sites for EB1 (16), and an array of binding sites for other proteins including p150 (reviewed by (3)). However, while it seems widely known that CLIP-170 contains IDRs, we were not able to find a formal analysis of this question in the literature.

Therefore, we evaluated the sequence of human CLIP-170 for the presence of disordered regions by using the established predictors “ESpritz version 1.3” (50), “MobiDB-lite” (51), and “IUPred” (52). In parallel, we used the “COILS” server to delineate the coiled-coil region (Fig 6) because we wanted to focus on IDR predictions outside of likely coiled-coil; this was important because in preliminary work, some predictors (e.g.,PONDR) frequently identified coiled-coil regions as likely IDRs (see also (61)). These analyses show that the three serine-rich regions in the N-terminal domain and regions adjacent to the zinc-knuckles in the C-terminal domain are predicted to be disordered in human CLIP-170 (Fig 6).

The observations that CLIP-170 is multivalent (3, 15, 16) and contains IDRs (Fig 6) are consistent with the idea that CLIP-170 can participate in biomolecular condensates. If these attributes are functionally significant, they should be conserved in diverse organisms. Indeed, examination of CLIP-170 relatives shows that the domain structure of CLIP-170 is conserved across diverse organisms (Fig 7 and S4 Fig), as previously noted in many publications; conservation of the domain structure implies conservation of multivalency. Moreover, our analysis demonstrates that the presence of IDRs adjacent to the CAP-Gly motifs and zinc knuckles is also well-conserved, extending even to the yeasts *S. cerevisiae* and *S. pombe* (Fig 7 and S4 Fig).

**Fig 7.**
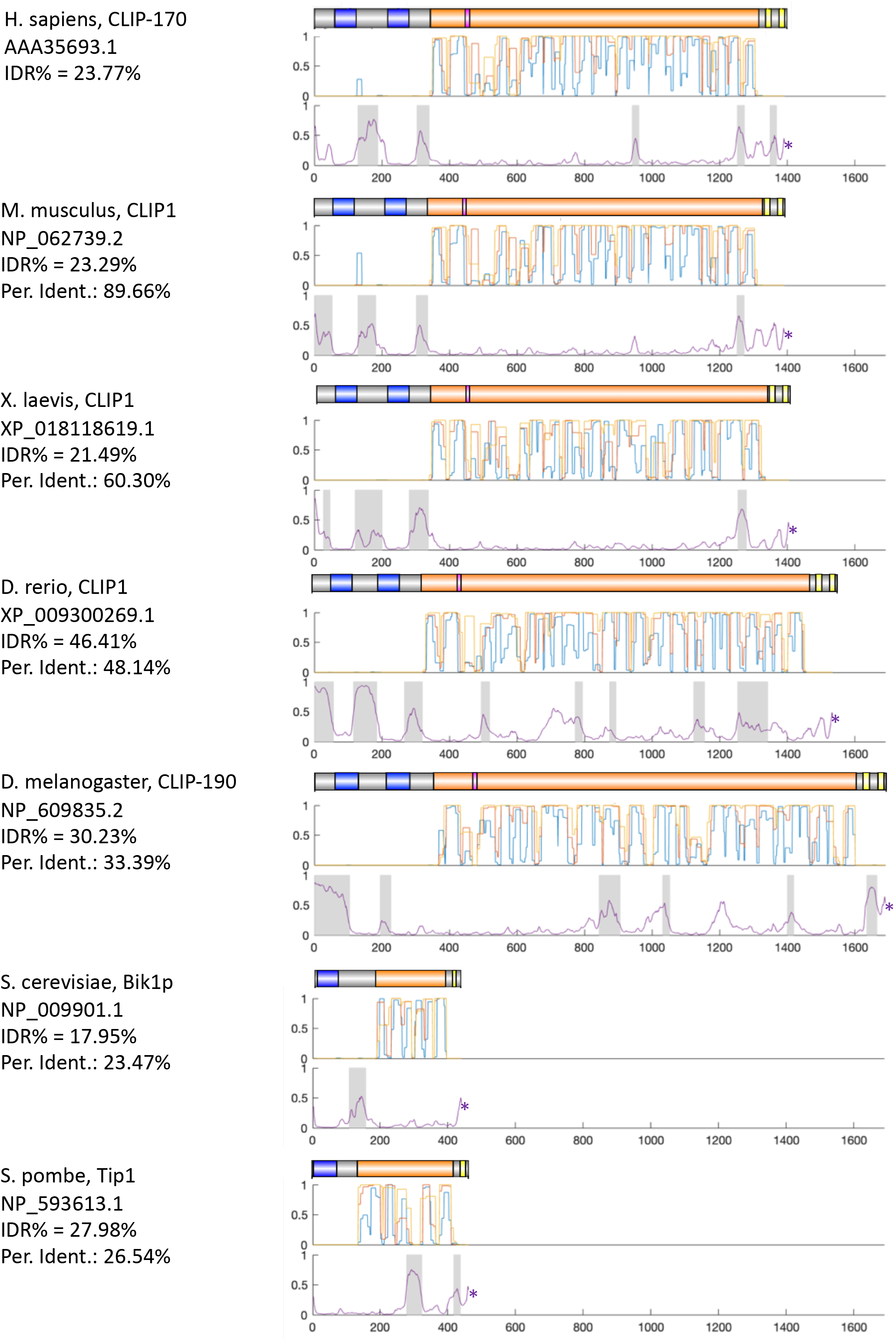
Analysis of coiled-coil and IDRs in CLIP-170 relatives from a range of different species. For each organism listed, one representative CLIP-170 sequence was selected for analysis, the results of which are shown in three images. The set of upper images provides the domain structure for each sequence, with colors as indicated: blue: CAP-Gly domains; orange: coiled-coil regions; pink: FEED domain; yellow: zinc-knuckle motifs. The sets of middle and lower images respectively provide the probabilities that a region is coiled-coil (as predicted by COILS) or IDR (as predicted by Espritz). For both analyses, 1 indicates 100% likelihood and * indicates the end of the sequence.

Two additional points are relevant to interpreting these data. First, though IDRs are not universally detected in the same places in all the organisms shown, assignment of IDRs is still an imperfect process; it remains possible that some of the proteins that appear to lack IDRs in particular regions (e.g., serine-rich 3 of Drosophila, the post-CAP-Gly serine-rich region of *S. pombe*) actually have them. Second, IDRs are sometimes identified in the coiled-coil regions, consistent with the established propensity of IDR prediction programs to find predicted IDRs in coiled-coil (61). The observation that these predicted IDRs are less conserved in their positions than are the IDRs near the CAP-Gly domains and zinc knuckles suggests that these may be spurious assignments.

In conclusion, the sequence characteristics of CLIP-170 and its relatives are consistent with the idea that CLIP-170 participates in biomolecular condensates and that this activity is conserved across a wide range of organisms.

### IDRs in other +TIP proteins are both common and conserved

Thus far, our analyses have focused on CLIP-170 and the condensates induced by CLIP-170 overexpression. As discussed in the Introduction, we are interested in the possibility that these large induced condensates reflect a much smaller +TIP network structure that exists under normal conditions, on the MT +TIP and/or in other cellular locations. Our sequence analysis (Fig 7 and S4 Fig) suggests that the ability of CLIP-170 to participate in condensates is likely to be conserved from humans to yeast, consistent with the idea that condensate formation is functionally significant. However, CLIP-170 can be deleted from some organisms (e.g., *S. cerevisiae*) with relatively minor effects (S1 Table), and other organisms (e.g., *Dictyostelium* amobebas, plants) lack CLIP-170 entirely, arguing against a central role for CLIP-170 in function of the +TIP network.

These observations indicate that if the normal activities of the +TIP network involve a functionally significant biomolecular condensate, other +TIPs should be able to drive condensate formation. Addressing this question in depth is beyond the scope of this paper. However, as a first step, we examined the sequences of other +TIPs to see if they have characteristics consistent with condensate formation, i.e., multivalency and the presence of IDRs. As noted above, it is already well-established that many human +TIPs are multivalent (3). Therefore, we examined the sequences of other human +TIPs to see if they contain IDRs. As part of this analysis, we included other proteins that colocalized with CLIP-170 from our immunofluorescence study (Fig 8).

**Fig 8.**
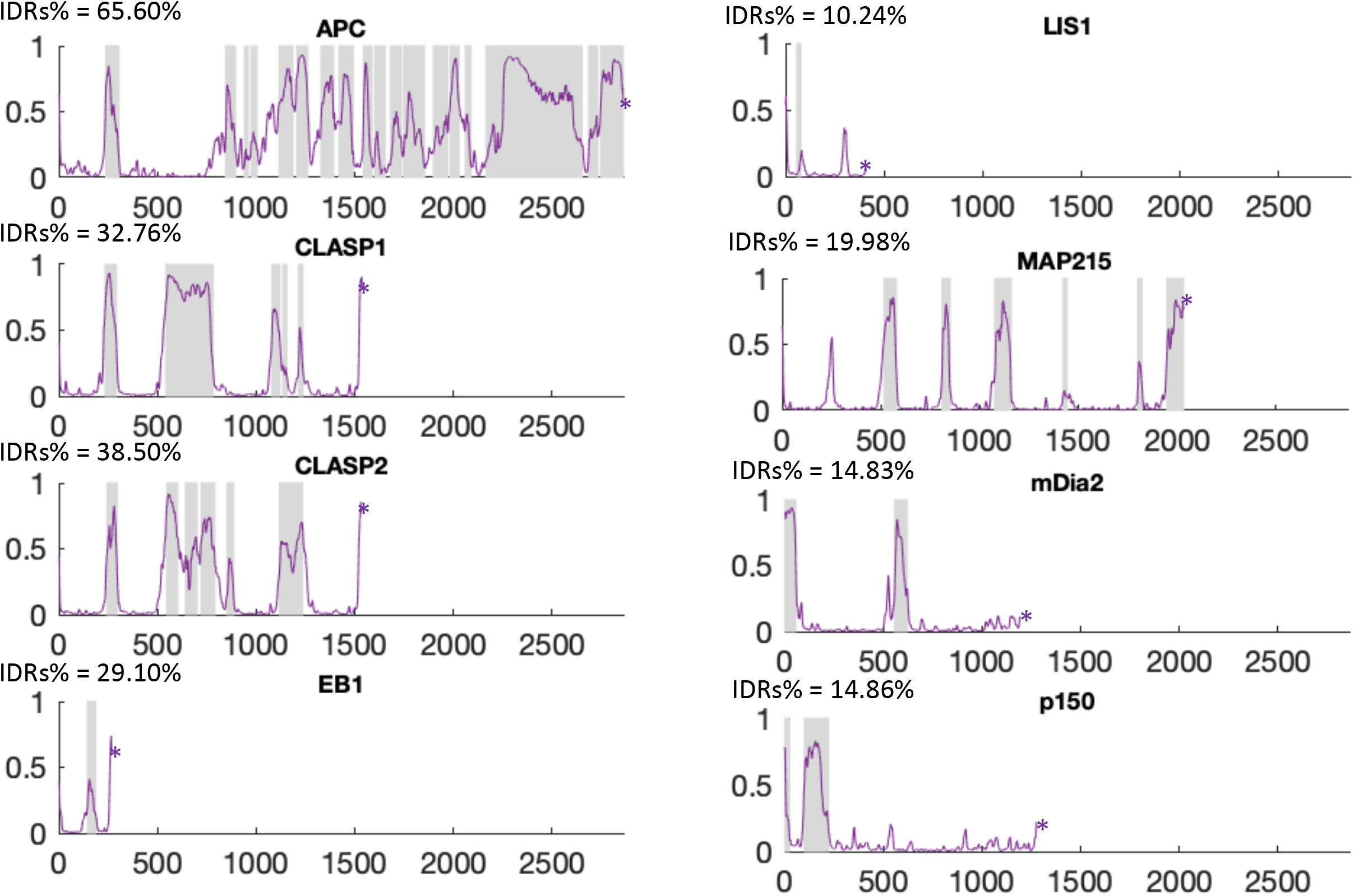
Analyses of IDRs in members of the human +TIP network. The Y-axis provides the probability of IDRs as a function of amino acid position (X-axis); 1 indicates 100% likelihood. The purple line provides the data for Espritz; shaded areas are the disordered regions predicted by MobiDB-lite. See Materials and Methods for the accession numbers.

Our analysis shows that the human versions of proteins that colocalized with CLIP-170 patches (EB1, LIS1, mDia2, and p150) contain predicted IDRs, as do other common +TIPs (APC, CLASP1, CLASP2, and MAP215). As part of this work, we also examined sequences for the presence of alpha helical coiled-coil (using the COILS server) and the other conserved domains (using hmmscan to search the PFAM database). Comparison of these results with the IDR predictions shows that the predicted disordered regions are mostly outside of alpha-helical coiled-coil and recognized folded domains (e.g. the CH domain in EB1) (S5 Fig), consistent with the idea that the IDR predictions are accurate. Although prediction of IDRs from sequence information is still maturing, and though presence of IDRs in a protein does not necessarily indicate that the protein participates in condensate formation, our results show that IDRs are very common in proteins involved in the +TIP network.

If the IDRs are functionally significant, their presence and perhaps also their positions relative to other conserved domains should be conserved across divergent organisms. To test these predictions, we chose the three most ubiquitous and conserved +TIPs (EB1, MAP215, and CLASP) and examined their sequences for the presence of IDRs and recognized domains in organisms ranging from humans to plants. The results (S6-S8 Figs) show that both the presence and position of disordered regions is well-conserved in all three proteins throughout all of the species examined.

More specifically, in EB1 (S6 Fig), both MobiDB-lite and Espritz predict a disordered region for all species upstream of the coiled-coiled region, between the CH and EBH domains. The Espritz predictor also predicts a disordered region after the EBH domain for all species (S6 Fig). In MAP215, both predictors find disordered regions outside of the TOG regions in all species. The pattern of disordered regions is surprisingly similar for all species, except in yeast, which is altered because of the reduced length of the MAP215 homologs in these organisms (S7 Fig). In CLASP, the presence and position of disordered regions are highly conserved from human to fly. The similarity of the domain structures is less obvious in yeast, *Dictyostelium*, and plants, but it is notable that the CLASP homologs in all of these organisms still contain IDRs (S8 Fig).

In conclusion, analysis of the protein sequences demonstrates that CLIP-170 and other members of the +TIP network have IDRs, consistent with participation of these proteins in biomolecular condensates. Moreover, the presence and position of these IDRs is well-conserved across animals and (where relevant) to organisms as divergent as plants, indicating that these IDRs are functionally significant. Moreover, the observation that the general domain structure of key +TIPs is conserved in organisms as diverse as plants (Fig 7 and S6-8 Figs) indicates that multivalency is also likely to be a conserved feature. These observations are consistent with the hypothesis that the +TIP network can assemble into a functionally significant biomolecular condensate.

## Discussion

### The “patches” induced by CLIP-170 are biomolecular condensates

Our results demonstrate that the so-called “patches” induced by CLIP-170 overexpression (8, 19) have the key hallmarks of biomolecular condensates (i.e., structures formed by liquid-liquid phase separation): elastic deformability (Fig 3 and Movie 1), the ability to undergo fission and fusion (Fig 3 and Movie 1), selective inclusion of some proteins and exclusions of others (Fig 4, S2 Fig, and Table 1), and rapid protein exchange with the cytoplasm (Fig 5). These observations lead us to conclude that the patches are biomolecular condensates. Significantly, the lack of overlap with established condensate markers (Fig 4, S2 Fig and S1 Fig) also suggests that the patches may represent a previously unrecognized type of condensate.

While these observations indicate that the CLIP-170-induced structures are biomolecular condensates, they do not address the question of functional significance. One possibility is that the overexpression-induced +TIP condensates are simply overexpression artifacts and so have no functional significance. Alternatively, the +TIP condensates observed in overexpression experiments could reflect +TIP superstructures that are formed by the endogenous +TIP network.

Addressing this question in a direct way would be challenging, in part because the comet at the end of MTs in unperturbed cells is both small (near the diffraction limit in size) and very dynamic. Moreover, a functionally significant +TIP condensate could form in environments other than the plus-end comet (e.g., at the cell cortex). For these reasons, directly testing the hypothesis that +TIP condensate formation is functionally relevant is beyond the scope of this work. However, as a first step in this direction, we examined CLIP-170 and other +TIPs for the presence of sequence characteristics (specifically multivalency and the presence of IDRs) consistent with participation in condensates. We then investigated how conserved these characteristics are across evolution. Finding out whether condensate associated sequence characteristics are common in human +TIP network proteins was intended to answer the question of whether condensate participation is a general feature +TIP network proteins or more specific to CLIP-170. The conservation analysis was performed as a test of functional significance, since a feature that impacts function of a basic cell biological process would be expected to be maintained across evolution.

As known from previous work (3, 15, 16), multivalency is indeed common in +TIPs (see also Fig 7 and S6-S8 Figs). Importantly, conservation of the basic domain structure of these proteins across wide spans of evolution (Fig 7 and S6-S8 Figs) indicates that multivalency is also a conserved feature of these proteins. In addition, we observed that IDRs are common in +TIPs (Fig 8 and S5 Fig). Significantly, for key +TIPs that can be recognized in divergent organisms, the existence and position of these predicted IDRs is also well-conserved across wide spans of evolution (e.g. humans through plants, Fig 7 and S6-S8 Figs), suggesting that these domains have activities that are functionally significant.

In considering these observations, it is important to stress that it is the existence and position of the predicted IDRs relative to conserved domains that are conserved, not the linear sequence of these predicted IDR-domains themselves. Indeed, at a primary sequence level, the regions identified as IDRs typically align poorly, once organisms outside of animals are compared (e.g., S4 Fig). Although one might be tempted to conclude that the poor alignability of these regions suggests lack of functional significance, it is instead consistent with the idea that these regions of the +TIPs are intrinsically disordered (62). In conclusion, our sequence analysis indicates that multiple members of the +TIP network have sequence features consistent with condensate formation and that these features are largely conserved across evolution, providing evidence that +TIP condensate formation is functionally significant.

So, starting from the recognition that we do not yet know whether +TIP condensate formation is physiologically significant, how *might* it be physiologically significant? As discussed more below, we propose that condensate formation may be fundamental to the core function of the +TIP network in regulating MT dynamics. As a foundation for discussing this hypothesis, we first review established functions for condensates and briefly review what is known about the mechanism of MT dynamic instability and its regulation by MT binding proteins.

### Functions of established biomolecular condensates

Before considering how a +TIP condensate might function, it is useful to first examine functions of other condensates. As discussed in the Introduction, the function of condensates is typically less clear than their existence. However, two commonly proposed functions are concentrating biomolecules to promote reactions (e.g., (37)) and segregating biomolecules into storage depots to prevent unwanted reactions (e.g, (25)). Both functions involve using condensates to alter the concentrations of reactants and thus the rates or timing of reactions. A very different set of proposed functions involves the ability of condensates to resist mechanical force (38) or even generate it (63). Though sometime left of our discussions of condensate function, is not surprising that assembly of condensates (which could be considered a type of 3D polymerization) would be able to generate mechanical force, given the well-demonstrated ability of actin and tubulin polymerization to generate force (64, 65). Finally, another set of proposed functions for condensates, one which overlaps with the ideas above, is that they play a role in signal transduction (66). Indeed, one could imagine that a phase separated droplet, a local region of high concentration that is in communication with the rest of the cell, could be an ideal structure for integrating information from multiple pathways, helping the cell to produce a coherent response to conflicting signals (67).

While it is now well-known that condensates are important for processes such as stress response and RNA processing, recent work also indicates that condensate formation plays a fundamental role in cytoskeletal processes. For example, phase separation appears to play an important role in regulation of actin nucleation (68–70). In the MT cytoskeleton, there is evidence that condensates are important to both the normal function of the MT binding protein Tau and its role in neurodegenerative disease (34–36). Moreover, one of the most striking existing examples of functionally significant cytoplasmic condensates is the centrosome, where condensate formation helps to both concentrate and localize the microtubule nucleating machinery and resist the forces generated during cell division (38). Here we consider possible roles for a condensate composed of +TIP proteins.

### Mechanism of MT dynamic instability and its regulation by MT binding proteins

To consider possible functions for +TIP condensates and how they might regulate MT dynamic instability (DI), it is first necessary to consider the mechanism of DI. MT growth and shrinkage result from the incorporation and detachment of tubulin dimers at protofilament ends exposed on MT tips. The details of the mechanism of DI are still being elucidated, but models center around the observation that tubulin subunits bind GTP but hydrolyze it soon after hydrolysis; this and other work have led to the idea that MTs can grow as long as they maintain a cap of GTP tubulin but switch to depolymerization (i.e., undergo “catastrophe”) if they lose this cap through stochastic GTP hydrolysis or other mechanisms (reviewed by (2).

Additional detail in the process is provided by recent electron microscopy indicating that growing MTs have extended protofilaments that are laterally unbound near the tip (71). This observation suggests that protofilaments need to “zip up” to be fully incorporated into the MT, and thus that the tip might be both fragile and very dynamic. Though surprising to some, the existence of laterally unbound protofilaments was predicted by a dimer scale computational model (72), and it is consistent with biophysical experiments that indicate that subunits exchange rapidly at the tip (73).

An important and puzzling observation is that MTs *in vivo* grow ~5x faster than those *in vitro*, even when tubulin concentrations have been matched (74). Remarkably, mixtures of +TIPs (specifically EB1 and MAP215) can reconstitute *in vivo* rates of polymerization *in vitro* (7). Two explanations have been offered for how the proteins are able to enhance the growth rates so much. One possibility that is that MAP215 and its relatives enhance the addition of new tubulin subunits to the tip, e.g., by binding and “delivering” tubulin subunits (75). The other possibility is that the proteins increase the rate of growth by reducing the fraction tubulin subunits that detach after binding (4, 76).

Though less intuitive, this second mechanism is attractive as an explanation for the dramatic effect of +TIPs, especially when combined with evidence that MT tips have laterally unbound protofilaments (71). Briefly, the observation mentioned above that incoming tubulin subunits “exchange rapidly at the tip” means that most incoming subunit detach instead of becoming incorporated into the lattice (73). A protein that could prevent detachment could dramatically increase the rate of polymerization (76). Laterally unbound protofilaments as discussed above might be expected to break off in chunks. Thus, one straightforward way for +TIPs enhance the net growth rate so dramatically would be to prevent detachment by zipping up the bonds between protofilaments (4).

### Hypothesis: the +TIP network forms a condensate that coats the MT tip, creating a superstructure that acts as a polymerization chaperone

As mentioned above, our lab has previously proposed that the web of +TIP interactions may have a physical purpose, creating a superstructure that promotes MT polymerization by stabilizing the weak lateral interactions between protofilaments, thus increasing the fraction of incoming subunits that are incorporated into the lattice (4). Here we extend this hypothesis by proposing that +TIP network superstructure referred to in this original hypothesis is a biomolecular condensate. More specifically, we suggest that the endogenous +TIP network can form a stocking-like structure that acts as a polymerization chaperone by encircling the MT tip in a supportive web, increasing the rate of lateral bond formation between protofilaments, which in turn decreases the rate of subunit detachment and thus increases the overall rate of MT growth (Fig 9).

**Fig 9.**
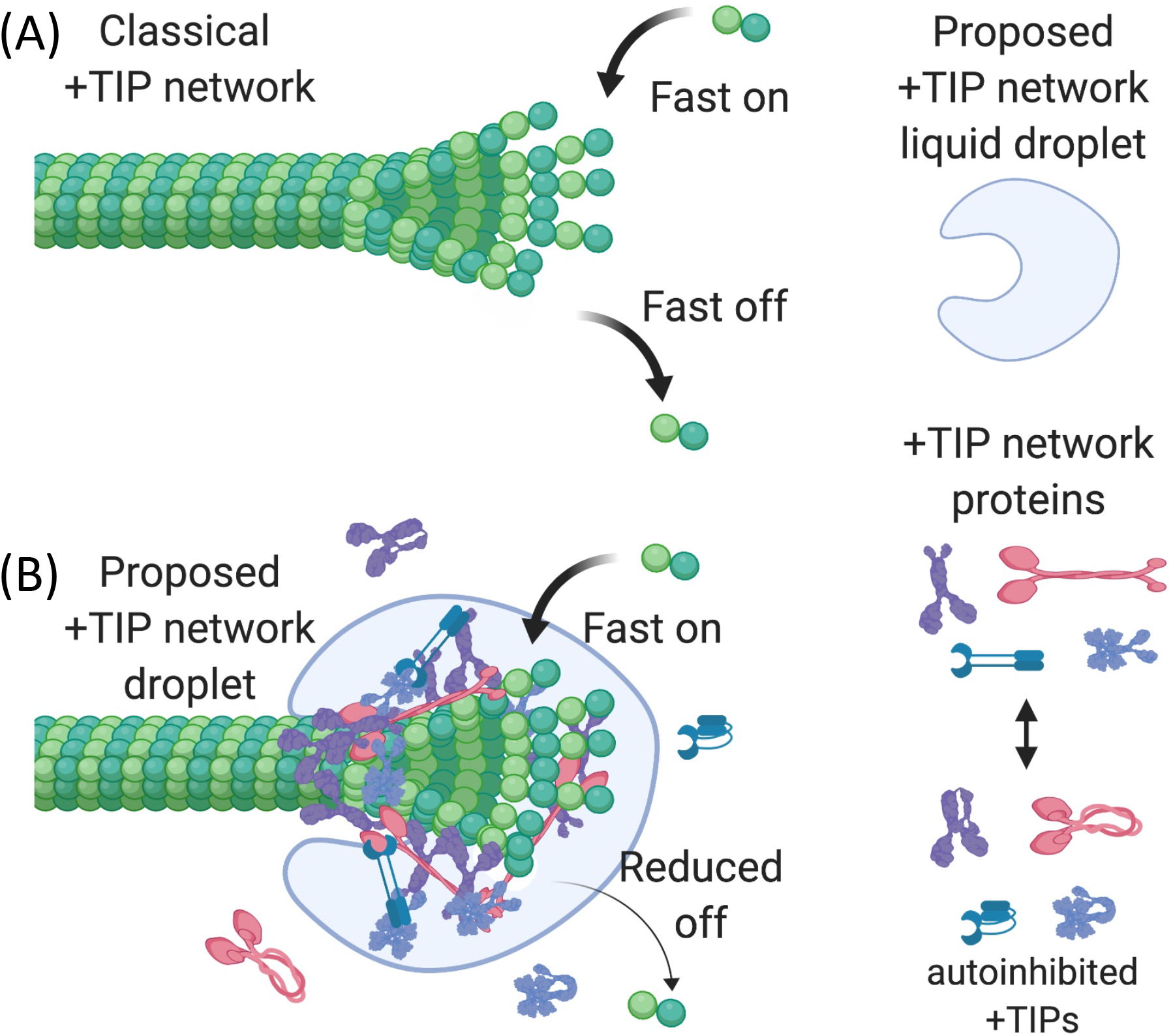
Proposed polymerization chaperone model. (A) In absence of the +TIP network, tubulin subunits arrive and leave quickly because most newly attached subunits are initially in the laterally unbonded regions of the tip protofilaments. The result is that MTs grow relatively slowly because most subunits detach before being incorporated into the laterally bound region of the MT. (B) In the presence of the proposed +TIP network liquid droplet, the web of interaction between the +TIPs creates a dynamic stocking-like structure that tracks the tip by diffusing on the potential energy gradient created by the preferential affinity of some tip components (most notably EB1) for tip-specific tubulin conformations. As the droplet moves, it exerts a force on the tip that promotes the zipping up of lateral bonds between protofilaments. This effect in turn increases the likelihood that newly arrived subunits are incorporated into the lattice and enables the MT to grow faster, as seen *in vivo* or with mixtures of +TIPs *in vitro* (4). This figure was created with BioRender (5).

As indicated in Fig 9, such a structure could potentially track the growing end by diffusing on the “wave” formed by the GTP (or GDP-Pi) cap, e.g., as has been proposed for proteins such as XMAP215 (77). Alternatively, one could also imagine +TIP droplet superstructure having utility if placed in a specific static position on a MT, such as has been suggested for Tau condensates (36).

Whether +TIP comets necessarily correspond to condensates is not specified in our hypothesis. Indeed, one piece of evidence against the idea that all +TIPs comets involve condensates is that researchers performing FRAP of MT comets using diffraction-limited spots measured a t_1/2_ of ~0.2 sec (78). This number is ~40 times faster than the t_1/2_ we observed for the CLIP-170 induced condensates (~8.5 sec). Regardless, even if the +TIP network on the average MT tip is not a condensate, one might expect that condensate formation could be triggered locally from +TIP localized proteins in response to appropriate signals. One locale where the possibility of +TIP condensate formation seems particularly intriguing is at the kinetochore-MT attachment site, given that a number of +TIP proteins localize to kinetochores and that condensate formation at the kinetochore has already been suggested (79). Indeed, the evidence cited above that condensates can generate mechanical force (63) is particularly striking when considered in the context of evidence that pulling forces on kinetochore components can promote MT growth (80, 81).

## Conclusion

In summary, we have shown that “patches” induced by CLIP-170 overexpression are biomolecular condensates. The overexpression approach, although artificial, may still give us insight to the mechanism of the physiological +TIP network. Initial support for the idea that condensate formation is functionally significant is provided by our bioinformatics studies, which have shown that CLIP-170 and other members of the +TIP network have conserved IDRs. This evidence leads us to propose that the endogenous +TIP network can under appropriate conditions form a dynamic stocking-like condensate that zips up the lateral bonds between protofilaments to promote MT polymerization.

## Acknowledgements

We are grateful to Optical Microscopy Core/NDIIF, University of Notre Dame, for the use of their instruments and services in support of this work. This research was funded by a fellowship from the American Heart Association (#17PRE33670896) to YOW and a grant from the National Science Foundation (MCB #1817966) to HVG.

## Reference

1. Matamoros AJ, Baas PW. Microtubules in health and degenerative disease of the nervous system. Brain Res Bull. 2016;126(Pt 3):217–25.

2. Goodson HV, Jonasson EM. Microtubules and Microtubule-Associated Proteins. Cold Spring Harb Perspect Biol. 2018;10(6).

3. Akhmanova A, Steinmetz MO. Control of microtubule organization and dynamics: two ends in the limelight. Nat Rev Mol Cell Biol. 2015;16(12):711–26.

4. Gupta KK, Alberico EO, Nathke IS, Goodson HV. Promoting microtubule assembly: A hypothesis for the functional significance of the +TIP network. Bioessays. 2014;36(9):818–26.

5. BioRender. 2020. Available from: https://biorender.com/.

6. Li W, Moriwaki T, Tani T, Watanabe T, Kaibuchi K, Goshima G. Reconstitution of dynamic microtubules with Drosophila XMAP215, EB1, and Sentin. J Cell Biol. 2012;199(5):849–62.

7. Zanic M, Widlund PO, Hyman AA, Howard J. Synergy between XMAP215 and EB1 increases microtubule growth rates to physiological levels. Nat Cell Biol. 2013;15(6):688–93.

8. Pierre P, Scheel J, Rickard JE, Kreis TE. CLIP-170 links endocytic vesicles to microtubules. Cell. 1992;70(6):887–900.

9. Pierre P, Pepperkok R, Kreis TE. Molecular characterization of two functional domains of CLIP-170 in vivo. J Cell Sci. 1994;107 (Pt 7):1909–20.

10. Lomakin AJ, Kraikivski P, Semenova I, Ikeda K, Zaliapin I, Tirnauer JS, et al. Stimulation of the CLIP-170--dependent capture of membrane organelles by microtubules through fine tuning of microtubule assembly dynamics. Mol Biol Cell. 2011;22(21):4029–37.

11. Komarova YA, Akhmanova AS, Kojima S, Galjart N, Borisy GG. Cytoplasmic linker proteins promote microtubule rescue in vivo. J Cell Biol. 2002;159(4):589–99.

12. Henty-Ridilla JL, Rankova A, Eskin JA, Kenny K, Goode BL. Accelerated actin filament polymerization from microtubule plus ends. Science. 2016;352(6288):1004–9.

13. Lewkowicz E, Herit F, Le Clainche C, Bourdoncle P, Perez F, Niedergang F. The microtubule-binding protein CLIP-170 coordinates mDia1 and actin reorganization during CR3-mediated phagocytosis. J Cell Biol. 2008;183(7):1287–98.

14. Noritake J, Watanabe T, Sato K, Wang S, Kaibuchi K. IQGAP1: a key regulator of adhesion and migration. J Cell Sci. 2005;118(Pt 10):2085–92.

15. Gupta KK, Paulson BA, Folker ES, Charlebois B, Hunt AJ, Goodson HV. Minimal plus-end tracking unit of the cytoplasmic linker protein CLIP-170. J Biol Chem. 2009;284(11):6735–42.

16. Chen Y, Wang P, Slep KC. Mapping multivalency in the CLIP-170-EB1 microtubule plus-end complex. J Biol Chem. 2019;294(3):918–31.

17. Scheel J, Pierre P, Rickard JE, Diamantopoulos GS, Valetti C, van der Goot FG, et al. Purification and analysis of authentic CLIP-170 and recombinant fragments. J Biol Chem. 1999;274(36):25883–91.

18. Mishima M, Maesaki R, Kasa M, Watanabe T, Fukata M, Kaibuchi K, et al. Structural basis for tubulin recognition by cytoplasmic linker protein 170 and its autoinhibition. Proc Natl Acad Sci U S A. 2007;104(25):10346–51.

19. Goodson HV, Skube SB, Stalder R, Valetti C, Kreis TE, Morrison EE, et al. CLIP-170 interacts with dynactin complex and the APC-binding protein EB1 by different mechanisms. Cell Motil Cytoskeleton. 2003;55(3):156–73.

20. Weisbrich A, Honnappa S, Jaussi R, Okhrimenko O, Frey D, Jelesarov I, et al. Structure-function relationship of CAP-Gly domains. Nat Struct Mol Biol. 2007;14(10):959–67.

21. Banani SF, Lee HO, Hyman AA, Rosen MK. Biomolecular condensates: organizers of cellular biochemistry. Nat Rev Mol Cell Biol. 2017;18(5):285–98.

22. Alberti S, Gladfelter A, Mittag T. Considerations and Challenges in Studying Liquid-Liquid Phase Separation and Biomolecular Condensates. Cell. 2019;176(3):419–34.

23. Bracha D, Walls MT, Brangwynne CP. Probing and engineering liquid-phase organelles. Nat Biotechnol. 2019;37(12):1435–45.

24. Ditlev JA, Case LB, Rosen MK. Who’s In and Who’s Out-Compositional Control of Biomolecular Condensates. J Mol Biol. 2018;430(23):4666–84.

25. Brangwynne CP, Eckmann CR, Courson DS, Rybarska A, Hoege C, Gharakhani J, et al. Germline P granules are liquid droplets that localize by controlled dissolution/condensation. Science. 2009;324(5935):1729–32.

26. Kulkarni M, Ozgur S, Stoecklin G. On track with P-bodies. Biochem Soc Trans. 2010;38(Pt 1):242–51.

27. Sear RP. Dishevelled: a protein that functions in living cells by phase separating. Soft Matter. 2007;3(6):680–4.

28. Scheer U, Hinssen H, Franke WW, Jockusch BM. Microinjection of actin-binding proteins and actin antibodies demonstrates involvement of nuclear actin in transcription of lampbrush chromosomes. Cell. 1984;39(1):111–22.

29. Gammons M, Bienz M. Multiprotein complexes governing Wnt signal transduction. Curr Opin Cell Biol. 2018;51:42–9.

30. Woodruff JB, Ferreira Gomes B, Widlund PO, Mahamid J, Honigmann A, Hyman AA. The Centrosome Is a Selective Condensate that Nucleates Microtubules by Concentrating Tubulin. Cell. 2017;169(6):1066–77 e10.

31. Rale MJ, Kadzik RS, Petry S. Phase Transitioning the Centrosome into a Microtubule Nucleator. Biochemistry. 2018;57(1):30–7.

32. Cioce M, Lamond AI. Cajal bodies: a long history of discovery. Annu Rev Cell Dev Biol. 2005;21:105–31.

33. Lallemand-Breitenbach V, de The H. PML nuclear bodies. Cold Spring Harb Perspect Biol. 2010;2(5):a000661.

34. Wegmann S, Eftekharzadeh B, Tepper K, Zoltowska KM, Bennett RE, Dujardin S, et al. Tau protein liquid-liquid phase separation can initiate tau aggregation. EMBO J. 2018;37(7).

35. Ambadipudi S, Biernat J, Riedel D, Mandelkow E, Zweckstetter M. Liquid-liquid phase separation of the microtubule-binding repeats of the Alzheimer-related protein Tau. Nat Commun. 2017;8(1):275.

36. Tan R, Lam AJ, Tan T, Han J, Nowakowski DW, Vershinin M, et al. Microtubules gate tau condensation to spatially regulate microtubule functions. Nat Cell Biol. 2019;21(9):1078–85.

37. Snead WT, Gladfelter AS. The Control Centers of Biomolecular Phase Separation: How Membrane Surfaces, PTMs, and Active Processes Regulate Condensation. Mol Cell. 2019;76(2):295–305.

38. Mittasch M, Tran VM, Rios MU, Fritsch AW, Enos SJ, Ferreira Gomes B, et al. Regulated changes in material properties underlie centrosome disassembly during mitotic exit. J Cell Biol. 2020;219(4).

39. Choi JM, Holehouse AS, Pappu RV. Physical Principles Underlying the Complex Biology of Intracellular Phase Transitions. Annu Rev Biophys. 2020;49:107–33.

40. Diamantopoulos GS, Perez F, Goodson HV, Batelier G, Melki R, Kreis TE, et al. Dynamic localization of CLIP-170 to microtubule plus ends is coupled to microtubule assembly. J Cell Biol. 1999;144(1):99–112.

41. Perez F, Diamantopoulos GS, Stalder R, Kreis TE. CLIP-170 highlights growing microtubule ends in vivo. Cell. 1999;96(4):517–27.

42. Rickard JE, Kreis TE. Binding of pp170 to microtubules is regulated by phosphorylation. J Biol Chem. 1991;266(26):17597–605.

43. Simonsen A, Lippe R, Christoforidis S, Gaullier JM, Brech A, Callaghan J, et al. EEA1 links PI(3)K function to Rab5 regulation of endosome fusion. Nature. 1998;394(6692):494–8.

44. Kawano J, Ide S, Oinuma T, Suganuma T. A protein-specific monoclonal antibody to rat liver beta 1-->4 galactosyltransferase and its application to immunohistochemistry. J Histochem Cytochem. 1994;42(3):363–9.

45. Schindelin J, Arganda-Carreras I, Frise E, Kaynig V, Longair M, Pietzsch T, et al. Fiji: an open-source platform for biological-image analysis. Nat Methods. 2012;9(7):676–82.

46. Webster AW, M. An easy protocol for FRAP. 2014. Available from: http://www.its.caltech.edu/~bif/Public/An%20Easy%20Protocol%20for%20FRAP.docx.

47. Bulinski JC, Odde DJ, Howell BJ, Salmon TD, Waterman-Storer CM. Rapid dynamics of the microtubule binding of ensconsin in vivo. J Cell Sci. 2001;114(Pt 21):3885–97.

48. Sprague BL, Pego RL, Stavreva DA, McNally JG. Analysis of binding reactions by fluorescence recovery after photobleaching. Biophys J. 2004;86(6):3473–95.

49. Wachsmuth M. Molecular diffusion and binding analyzed with FRAP. Protoplasma. 2014;251(2):373–82.

50. Walsh I, Martin AJ, Di Domenico T, Tosatto SC. ESpritz: accurate and fast prediction of protein disorder. Bioinformatics. 2012;28(4):503–9.

51. Necci M, Piovesan D, Dosztanyi Z, Tosatto SCE. MobiDB-lite: fast and highly specific consensus prediction of intrinsic disorder in proteins. Bioinformatics. 2017;33(9):1402–4.

52. Meszaros B, Erdos G, Dosztanyi Z. IUPred2A: context-dependent prediction of protein disorder as a function of redox state and protein binding. Nucleic Acids Res. 2018;46(W1):W329–W37.

53. Erdos G, Dosztanyi Z. Analyzing Protein Disorder with IUPred2A. Curr Protoc Bioinformatics. 2020;70(1):e99.

54. Potenza E, Di Domenico T, Walsh I, Tosatto SC. MobiDB 2.0: an improved database of intrinsically disordered and mobile proteins. Nucleic Acids Res. 2015;43(Database issue):D315–20.

55. Sievers F, Wilm A, Dineen D, Gibson TJ, Karplus K, Li W, et al. Fast, scalable generation of high-quality protein multiple sequence alignments using Clustal Omega. Mol Syst Biol. 2011;7:539.

56. Mirarab S, Nguyen N, Guo S, Wang LS, Kim J, Warnow T. PASTA: Ultra-Large Multiple Sequence Alignment for Nucleotide and Amino-Acid Sequences. J Comput Biol. 2015;22(5):377–86.

57. Potter SC, Luciani A, Eddy SR, Park Y, Lopez R, Finn RD. HMMER web server: 2018 update. Nucleic Acids Res. 2018;46(W1):W200–W4.

58. Lupas A, Van Dyke M, Stock J. Predicting coiled coils from protein sequences. Science. 1991;252(5009):1162–4.

59. Bartolini F, Moseley JB, Schmoranzer J, Cassimeris L, Goode BL, Gundersen GG. The formin mDia2 stabilizes microtubules independently of its actin nucleation activity. J Cell Biol. 2008;181(3):523–36.

60. McSwiggen DT, Mir M, Darzacq X, Tjian R. Evaluating phase separation in live cells: diagnosis, caveats, and functional consequences. Genes Dev. 2019;33(23–24):1619–34.

61. van der Lee R, Buljan M, Lang B, Weatheritt RJ, Daughdrill GW, Dunker AK, et al. Classification of intrinsically disordered regions and proteins. Chem Rev. 2014;114(13):6589–631.

62. Moesa HA, Wakabayashi S, Nakai K, Patil A. Chemical composition is maintained in poorly conserved intrinsically disordered regions and suggests a means for their classification. Mol Biosyst. 2012;8(12):3262–73.

63. Wiegand T, Hyman AA. Drops and fibers - how biomolecular condensates and cytoskeletal filaments influence each other. Emerg Top Life Sci. 2020.

64. Simon C, Kusters R, Caorsi V, Allard A, Abou-Ghali M, Manzi J, et al. Actin dynamics drive cell-like membrane deformation. Nature Physics. 2019;15(6):602–9.

65. Brouhard GJ, Rice LM. Microtubule dynamics: an interplay of biochemistry and mechanics. Nat Rev Mol Cell Biol. 2018;19(7):451–63.

66. Case LB, Ditlev JA, Rosen MK. Regulation of Transmembrane Signaling by Phase Separation. Annu Rev Biophys. 2019;48:465–94.

67. Holehouse AS, Pappu RV. Functional Implications of Intracellular Phase Transitions. Biochemistry. 2018;57(17):2415–23.

68. Banjade S, Rosen MK. Phase transitions of multivalent proteins can promote clustering of membrane receptors. Elife. 2014;3.

69. Su X, Ditlev JA, Hui E, Xing W, Banjade S, Okrut J, et al. Phase separation of signaling molecules promotes T cell receptor signal transduction. Science. 2016;352(6285):595–9.

70. Case LB, Zhang X, Ditlev JA, Rosen MK. Stoichiometry controls activity of phase-separated clusters of actin signaling proteins. Science. 2019;363(6431):1093–7.

71. McIntosh JR, O’Toole E, Morgan G, Austin J, Ulyanov E, Ataullakhanov F, et al. Microtubules grow by the addition of bent guanosine triphosphate tubulin to the tips of curved protofilaments. J Cell Biol. 2018;217(8):2691–708.

72. Margolin G, Gregoretti IV, Cickovski TM, Li C, Shi W, Alber MS, et al. The mechanisms of microtubule catastrophe and rescue: implications from analysis of a dimer-scale computational model. Mol Biol Cell. 2012;23(4):642–56.

73. Gardner MK, Charlebois BD, Janosi IM, Howard J, Hunt AJ, Odde DJ. Rapid microtubule self-assembly kinetics. Cell. 2011;146(4):582–92.

74. Hyman AA, Karsenti E. Morphogenetic properties of microtubules and mitotic spindle assembly. Cell. 1996;84(3):401–10.

75. Al-Bassam J, Chang F. Regulation of microtubule dynamics by TOG-domain proteins XMAP215/Dis1 and CLASP. Trends Cell Biol. 2011;21(10):604–14.

76. Howard J, Hyman AA. Growth, fluctuation and switching at microtubule plus ends. Nat Rev Mol Cell Biol. 2009;10(8):569–74.

77. Brouhard GJ, Stear JH, Noetzel TL, Al-Bassam J, Kinoshita K, Harrison SC, et al. XMAP215 is a processive microtubule polymerase. Cell. 2008;132(1):79–88.

78. Dragestein KA, van Cappellen WA, van Haren J, Tsibidis GD, Akhmanova A, Knoch TA, et al. Dynamic behavior of GFP-CLIP-170 reveals fast protein turnover on microtubule plus ends. J Cell Biol. 2008;180(4):729–37.

79. Trivedi P, Palomba F, Niedzialkowska E, Digman MA, Gratton E, Stukenberg PT. The inner centromere is a biomolecular condensate scaffolded by the chromosomal passenger complex. Nat Cell Biol. 2019;21(9):1127–37.

80. Franck AD, Powers AF, Gestaut DR, Gonen T, Davis TN, Asbury CL. Tension applied through the Dam1 complex promotes microtubule elongation providing a direct mechanism for length control in mitosis. Nat Cell Biol. 2007;9(7):832–7.

81. Akiyoshi B, Sarangapani KK, Powers AF, Nelson CR, Reichow SL, Arellano-Santoyo H, et al. Tension directly stabilizes reconstituted kinetochore-microtubule attachments. Nature. 2010;468(7323):576–9.

