## Supplementary Material for "Overexpression of the microtubule-binding protein CLIP-170 induces a +TIP network superstructure consistent with a biomolecular condensate"

#### Supplement Table of Contents


| Figure/Table | Caption | Page |
|---|---|---|
| S1 Table | Example of +TIP knock-out mutants from literature | 2 |
| S1 Fig | Membrane colocalization studies suggest that CLIP-170 patches are membraneless structures | 3-4 |
| S2 Fig | CLIP-170 patches have selective properties | 5 |
| S3 Fig | Analysis of the size distribution of CLIP-170 condensates | 6 |
| S4 Fig | The position of CLIP-170 IDRs is conserved from yeast to humans | 7 |
| S5 Fig | Analysis of Coiled-coil domains and IDRs of +TIP network proteins | 8-9 |
| S6 Fig | Analysis of the position of coiled-coil domains and IDRs in EB1 across a range of organisms | 10-11 |
| S7 Fig | Analysis of the position of coiled-coil domains and IDRs in MAP215 across a range of organisms | 12-13 |
| S8 Fig | Analysis of the position of coiled-coil domains and IDRs in CLASPs across a range of organism | 14-16 |
| References | References for the supplementary material | 17-18 |


**Movie 1. Dynamic behaviors of CLIP-170 patches *in vivo*** (Corresponds to Fig 3). NIH3T3 cells were transfected to overexpress GFP-CLIP-170, and the behavior of GFP-CLIP-170 *in vivo* was recorded by confocal microscopy after 24-27 hr of transfection. Red box: An example of an apparent elastic deformation of a patch, following by fission. Arrows indicate examples of patch fusion (magenta) and a photobleached site (green).

#### Table S1. Example of +TIP knock-out mutants from literature

| Protein (family), organism | Paralogs present | Viability | Other phenotypes | Reference(s) |
|---|---|---|---|---|
| CLIP-170 (CLIP-170), mice | CLIP-115 | Viable | Male KO mice have abnormal sperm morphology. | (1, 2) |
| CLIP-115 (CLIP-170), mice | CLIP-170 | Viable | KO mice develop Williams syndrome, including mild growth deficiency and other neuron development related symptoms. | (2, 3) |
| CLIP-190 (CLIP-170), fruit fly | No | Viable | Flies with null allele of *clip-190* have no obvious defects. | (4) |
| BIK1 (CLIP-170), *S. cerevisiae* | No | Viable | Yeast with null-allele of *bik1* have a similar reproductive rate to wt, however, their nuclear positioning is abnormal. | (5-7) |
| EB1 (EB), HeLa cells | EB2, EB3 | Viable | Combined EB1/2/3 knock out HeLa cells can still go through mitosis but has some spindle abnormalities. | (8) |
| DmEB1 (EB), fruit fly | | Viable as larvae | Point mutation with reduced amount of DmEB1 is lethal due to failed eclosion. Surviving adults have abnormal wings and are flightless, which indicate neuromuscular defects. | (9) |
| BIM1 (EB), *S. cerevisiae* | No | Viable | Synthetic lethality with *bik1*, *num1*, and *bub3* deletion alleles. | (7, 10) |
| AtEB1a (EB), *Arabidopsis* | AtEB1b, AtEB1c | Viable | Single or multiple-deletion of EB1 homologous genes have mild phenotypes (skewed root). | (11) |
| Stu1 (CLASPs), *S. cerevisiae* | No | Inviable | Null mutant germinates and undergoes 1-2 cell divisions before cell division ceases. | (7, 12) |
| CLASP (CLASPs), *Arabidopsis* | No | Viable | Adult null mutant plants grow smaller, and have shorter inflorescences. | (13) |
| Stu2 (MAP215), *S. cerevisiae* | No | Inviable | Null mutant germinates and undergoes 3-4 cell divisions before cell division ceases. | (14) |
| Alp1 (MAP215), *S. pombe* | Dis1 | Viable | Δ*alp14* mutant is viable but has abnormal mitotic progression. Δ*alp14* mutant is synthetic lethal with *dis1* and *mad2*. | (15) |
| MACF1 (spectraplakin), mice | MACF2 | Inviable | Mutant mice are not viable because of neuronal migration defects. | (2) |
| MACF2 (spectraplakin), mice | MACF1 | Viable | KO mice develop severe dystonia and sensory nerve degeneration. | (2, 16) |

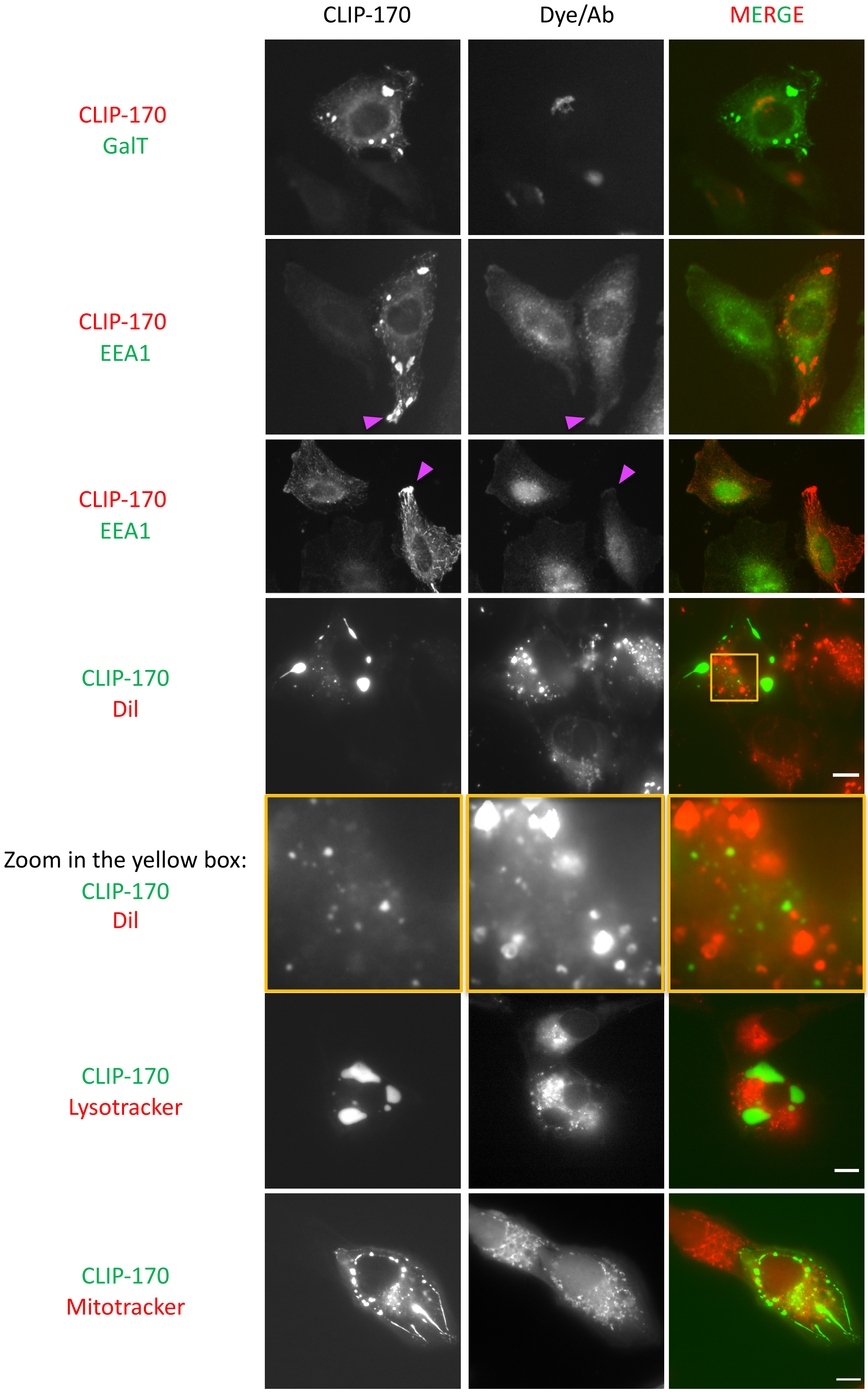
CLIP-170
Dye/Ab
MERGE
CLIP-170
GalT
CLIP-170
EEA1
CLIP-170
EEA1
CLIP-170
DiI
Zoom in the yellow box:
CLIP-170
DiI
CLIP-170
Lysotracker
CLIP-170
Mitotracker

**S1 Fig. Membrane colocalization studies suggest that CLIP-170 patches are membraneless structures.** For the EEA1 and GalT experiments, HeLa cells were transfected with full-length CLIP-170 for 24-36 hours, then fixed with methanol before immunofluorescence imaging with antibodies as specified; the magenta arrows indicate partial colocalization between EEA1 and CLIP-170 patches. For the other experiments, NIH3T3 cells were transfected with GFP-CLIP-170 for 24-25 hours, stained by the specified lipid or organelle markers, and imaged live by widefield microscopy. Scale bar: 10 µm.

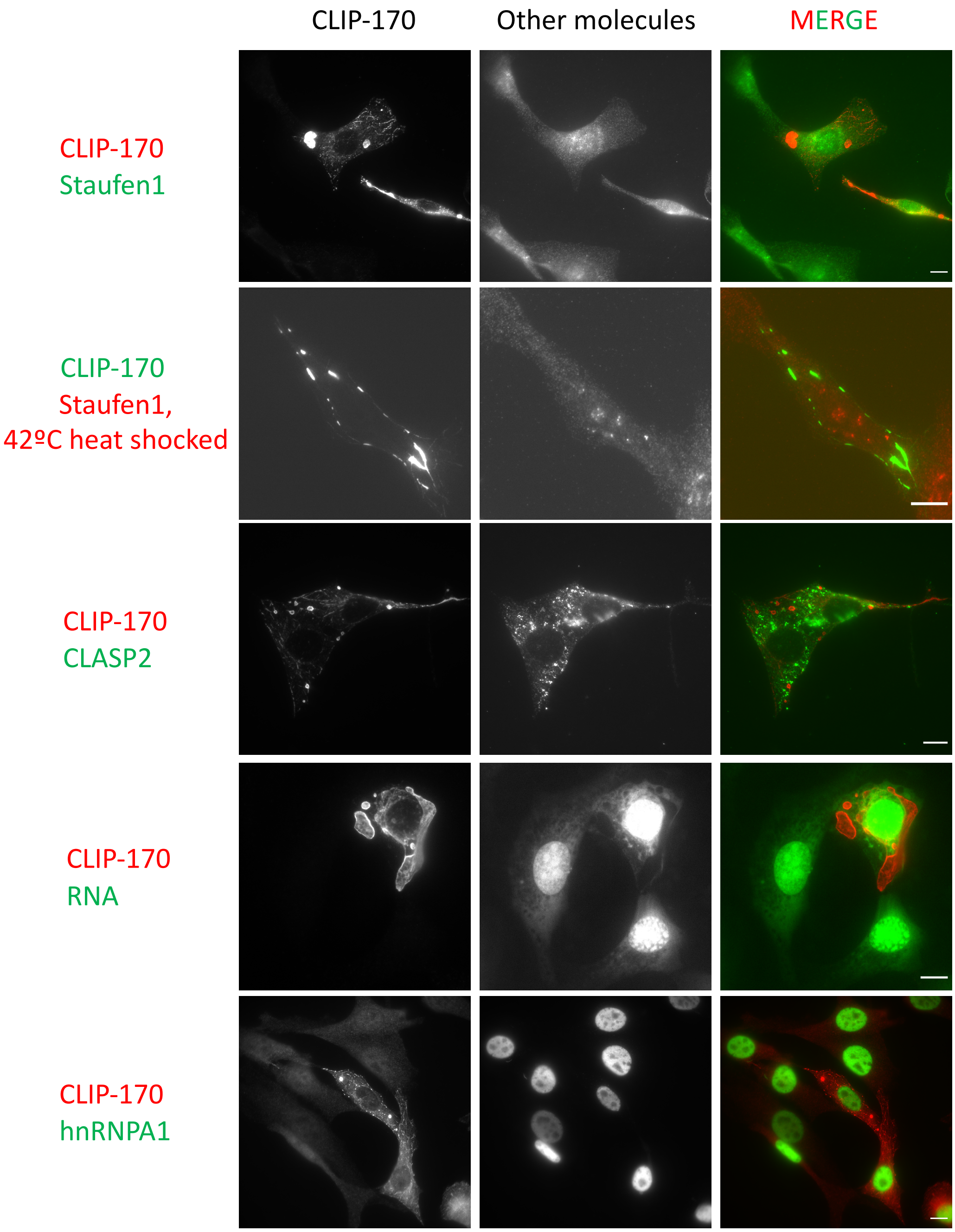


**S2 Fig. CLIP-170 patches have selective properties.** NIH3T3 cells were transfected with full-length CLIP-170 for 24 hr, fixed with PFA, probed with antibodies against molecules of interest (as indicated), and observed by widefield fluorescence microscopy. The contrast of each representative image in a given row is adjusted to the same levels. Scale bar: 10 µm.

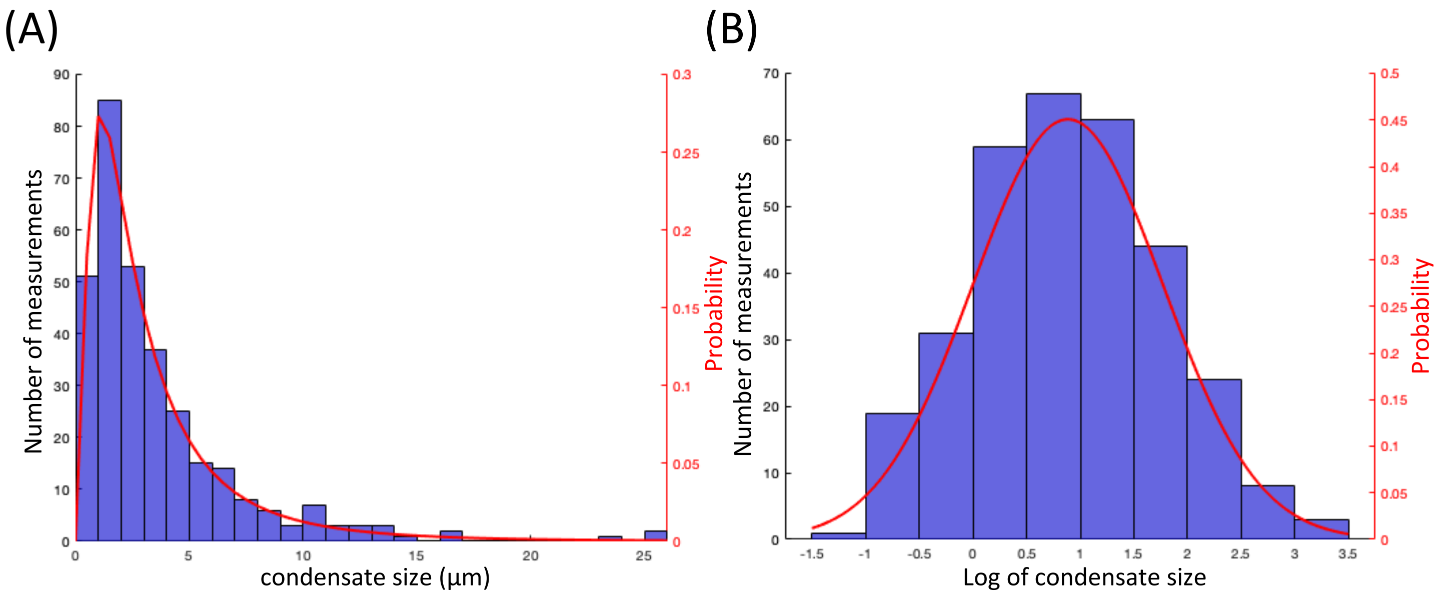


**S3 Fig. Analysis of the size distribution of CLIP-170 condensates**. To identify appropriate bin boundaries for separating condensates by size (Figs 5C-E), we used parametric distribution. (A) Histogram of condensate sizes. *Image acquisition and quantification:* NIH3T3 cells were transfected with GFP-CLIP-170. Cells were fixed with PFA after 24 hr of transfection, and 100 arbitrarily chosen fields of view were acquired using widefield fluorescence microscopy. For each condensate present in these images, we measured the widest width using Fiji, and this value was defined as the droplet size. A total of 319 condensates were measured. *Analysis of the distribution of condensate sizes:* The histogram of the condensate size distribution was plotted by the MATLAB "histogram" function. To determine the optimal bin size, we used the square-root model (a method commonly used for assessing skewed distributions). A Log-normal distribution was used to fit the histogram data. Finally, the "logninv" function was used to calculate the probability of condensate size falling within a certain range. We defined the smallest 50% of the condensates as small (0 - 2.4 µm), 50-75% as medium (2.4 - 4.4 µm), and >75% as large (≥ 4.4 µm). (B) Histogram of the log of condensate sizes. To test whether the log-normal distribution was an appropriate parametric distribution to use for these data, we took the log of condensate sizes and fit it with a normal distribution, using Sturge's rule to determine the optimal bin size. These results support our decision to use a log-normal fit because the plot shows that our raw data in (A) is distributed in a log normal form.

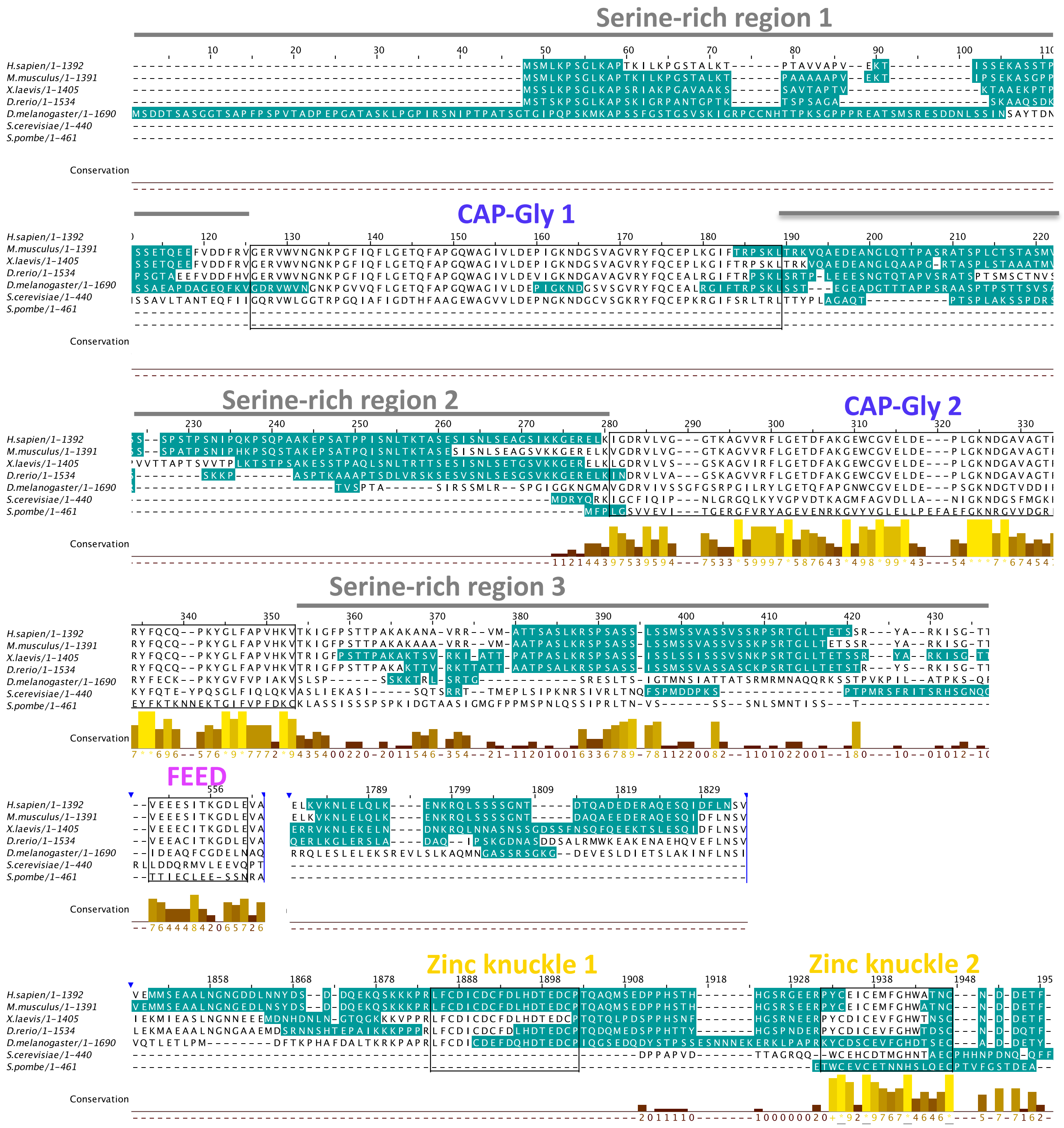


**S4 Fig. The position of CLIP-170 IDRs is conserved from yeast to humans.** Sequences here are the same as used in Fig 7. Cyan: disordered regions predicted by Espritz. Sequences are labeled with position in the alignment. Corresponding domain or motif structures were annotated above the aligned sequences. Annotations below were generated by Jalview and show the degree of conservation of the aligned columns.

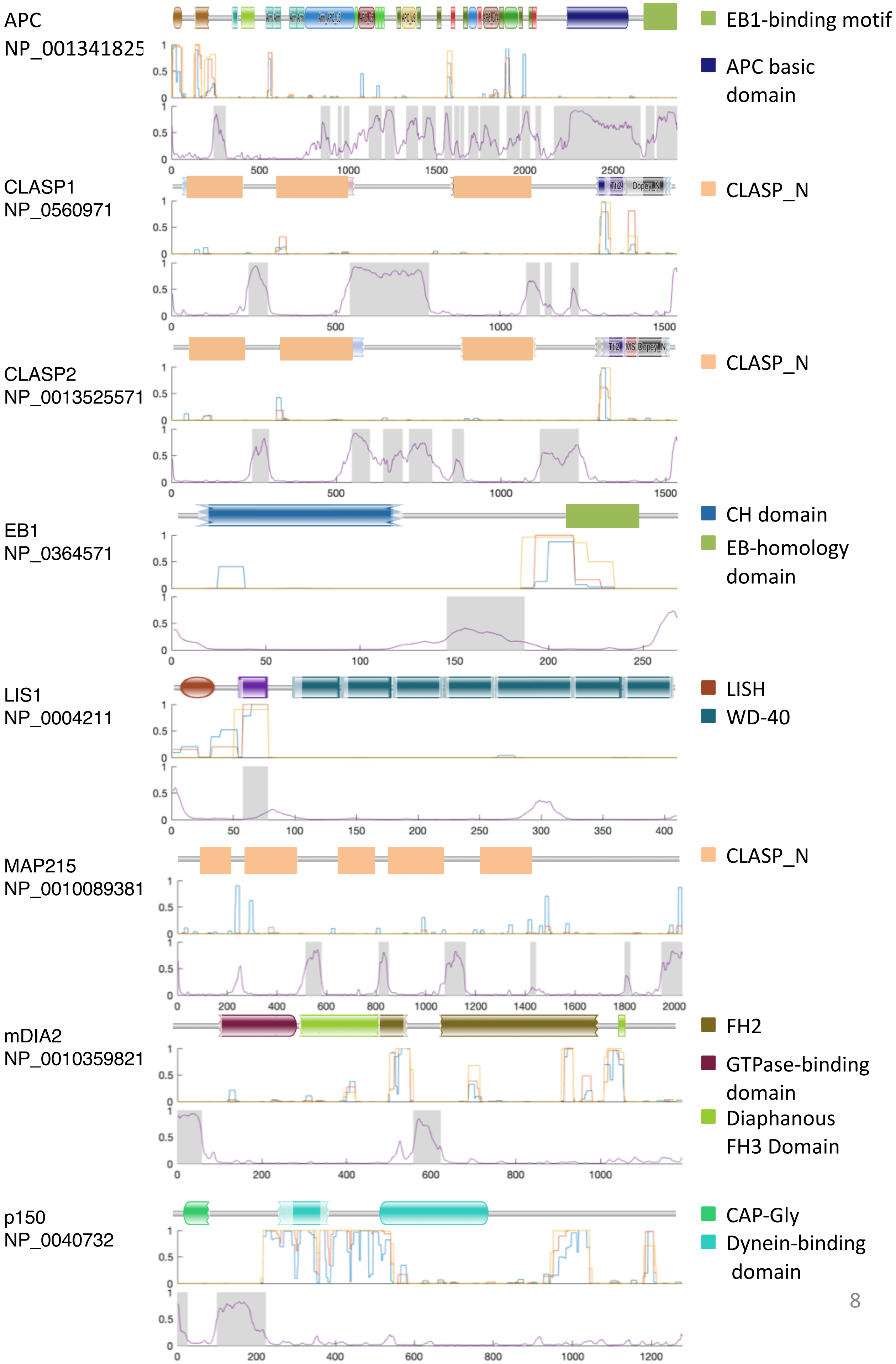

APC
NP_001341825
EB1-binding motif
APC basic domain
CLASP1
NP_0560971
CLASP_N
CLASP2
NP_0013525571
CLASP_N
EB1
NP_0364571
CH domain
EB-homology domain
LIS1
NP_0004211
LISH
WD-40
MAP215
NP_0010089381
CLASP_N
mDIA2
NP_0010359821
FH2
GTPase-binding domain
Diaphanous FH3 Domain
p150
NP_0040732
CAP-Gly
Dynein-binding domain

**S5 Fig. Analysis of Coiled-coil domains and IDRs of +TIP network proteins** (companion to Fig 8). For each +TIP network protein (accession number at left), the top image shows domain structures identified by Pfam (e-value < 0.1). For ease of visualization, we manually colored the most significant domains (as indicated on the right), while minor domains are shown as plotted by Pfam. Note that a few recognized domain structures were not predicted with the e-value we selected, meaning that the absence of a domain in this figure should not necessarily be interpreted as absence of the domain in the protein. The middle image represents the probability of coiled-coil region as predicted by COILS with three different windows (7AA, blue; 14 AA, orange; 21 AA yellow). In the bottom image, the purple line shows the probability of IDRs predicted by Espritz, and the shaded area indicates the IDRs as predicted by MobiDB-lite. For both COILS and Espritz analyses, 1 indicates 100% likelihood.

**Human EB1**

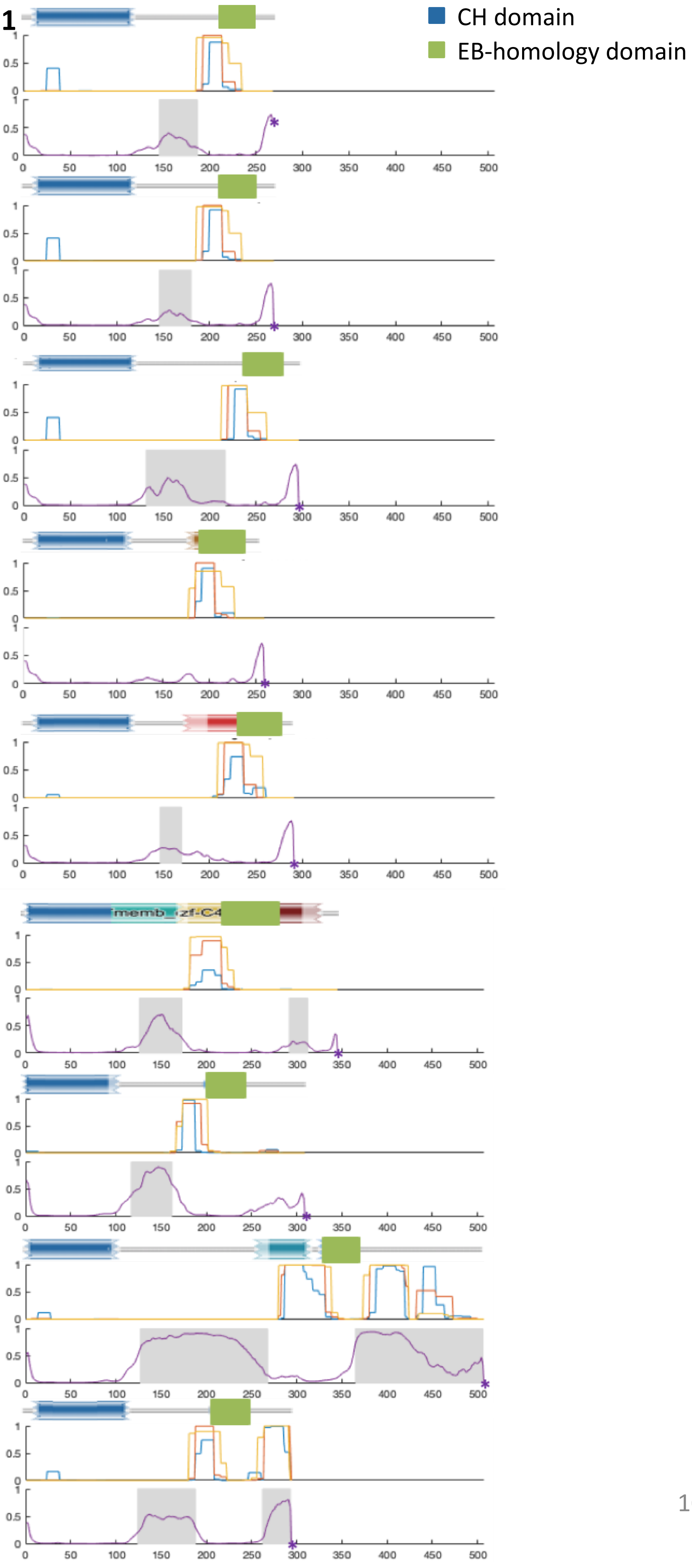


H. sapiens
NP_036457.1,
IDR% = 29.10%

M. musculus
NP_031922.1
IDR% = 20.89%

X. laevis
NP_001080896.1
IDR% = 26.10%

D. rerio
NP_998805.1
IDR% = 12.01%

D. melanogaster
NP_995752.1
IDR% = 27.93%

S. cerevisiae
NP_010932.1
IDR% = 25.87%

S. pombe
NP_593678.1
IDR% = 41.88%

D. discoidium
XP_638940.1
IDR% = 64.22%

A. thaliana
NP_201056.1
IDR% = 27.20%

**S6 Fig. Analysis of the position of coiled-coil domains and IDRs in EB1 across a range of organisms.** For each EB1 protein (one chosen per species, accession number at left), the top image shows the domain structures identified by Pfam (e-value < 0.1). For ease of visualization, we manually colored the most significant domains (as indicated on the right), while minor domains are shown as plotted by Pfam. Note that a few recognized domain structures were not predicted with the e-value we selected, meaning that the absence of a domain in this figure should not necessarily be interpreted as absence of the domain in the protein. The middle image represents the probability of coiled-coil region as predicted by COILS with three different windows (7AA, blue; 14 AA, orange; 21 AA yellow). In the bottom image, the purple line shows the probability of IDRs predicted by Espritz, and the shaded area indicates the IDRs as predicted by MobiDB-lite. For both COILS and Espritz analyses, 1 indicates 100% likelihood. * indicates the end of the sequence.

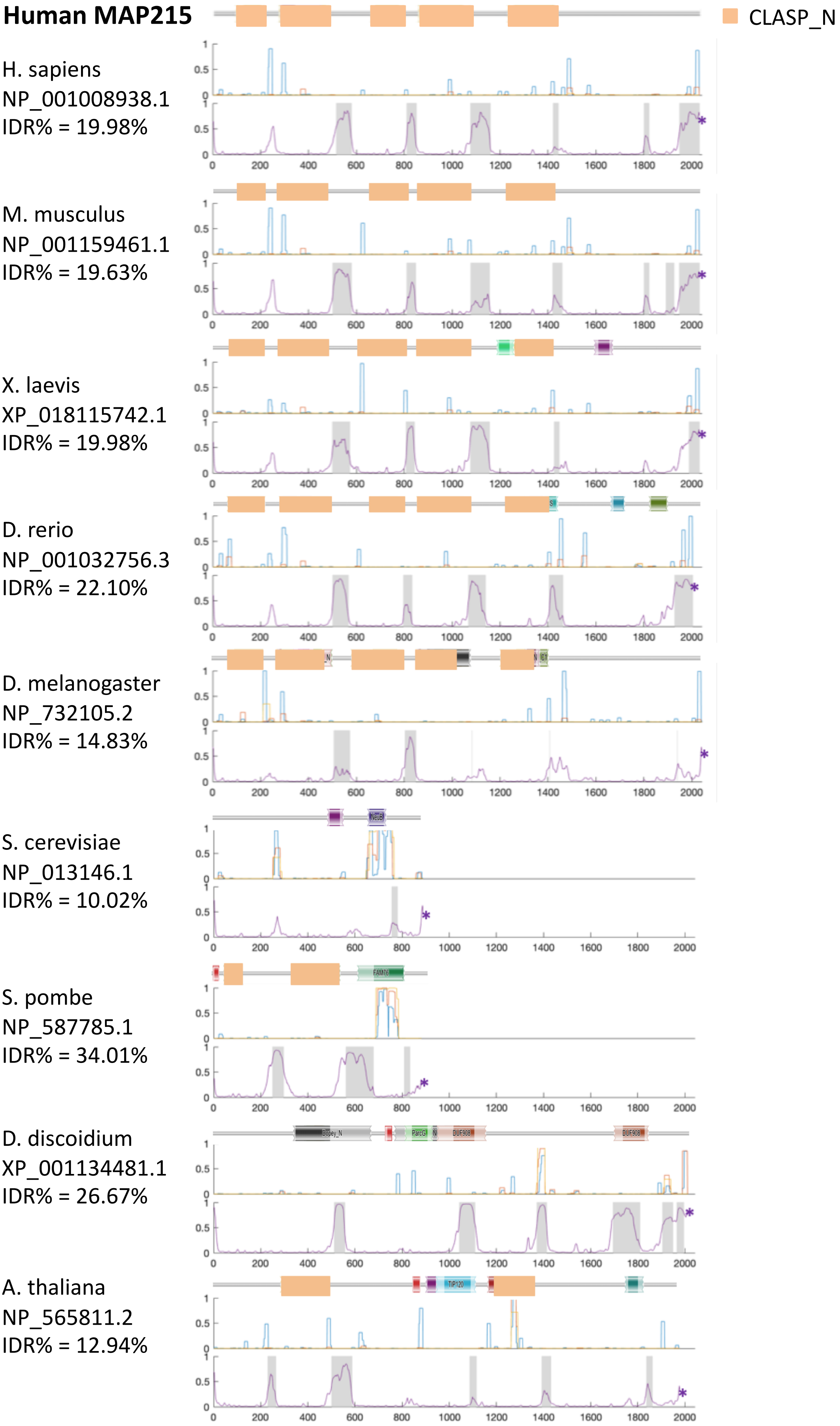
Human MAP215
CLASP_N
H. sapiens
NP_001008938.1
IDR% = 19.98%
M. musculus
NP_001159461.1
IDR% = 19.63%
X. laevis
XP_018115742.1
IDR% = 19.98%
D. rerio
NP_001032756.3
IDR% = 22.10%
D. melanogaster
NP_732105.2
IDR% = 14.83%
S. cerevisiae
NP_013146.1
IDR% = 10.02%
S. pombe
NP_587785.1
IDR% = 34.01%
D. discoidium
XP_001134481.1
IDR% = 26.67%
A. thaliana
NP_565811.2
IDR% = 12.94%

**S7 Fig. Analysis of the position of coiled-coil domains and IDRs in MAP215 across a range of organisms.** For each MAP215 protein (one chosen per species, accession number at left), the top image shows the domain structures identified by Pfam (e-value < 0.1). For ease of visualization, we manually colored the most significant domains (as indicated on the right), while minor domains are shown as plotted by Pfam. Note that a few recognized domain structures were not predicted with the e-value we selected, meaning that the absence of a domain in this figure should not necessarily be interpreted as absence of the domain in the protein. The middle image represents the probability of coiled-coil region as predicted by COILS with three different windows (7AA, blue; 14 AA, orange; 21 AA yellow). In the bottom image, the purple line shows the probability of IDRs predicted by Espritz, and the shaded area indicates the IDRs as predicted by MobiDB-lite. For both COILS and Espritz analyses, 1 indicates 100% likelihood. * indicates the end of the sequence.

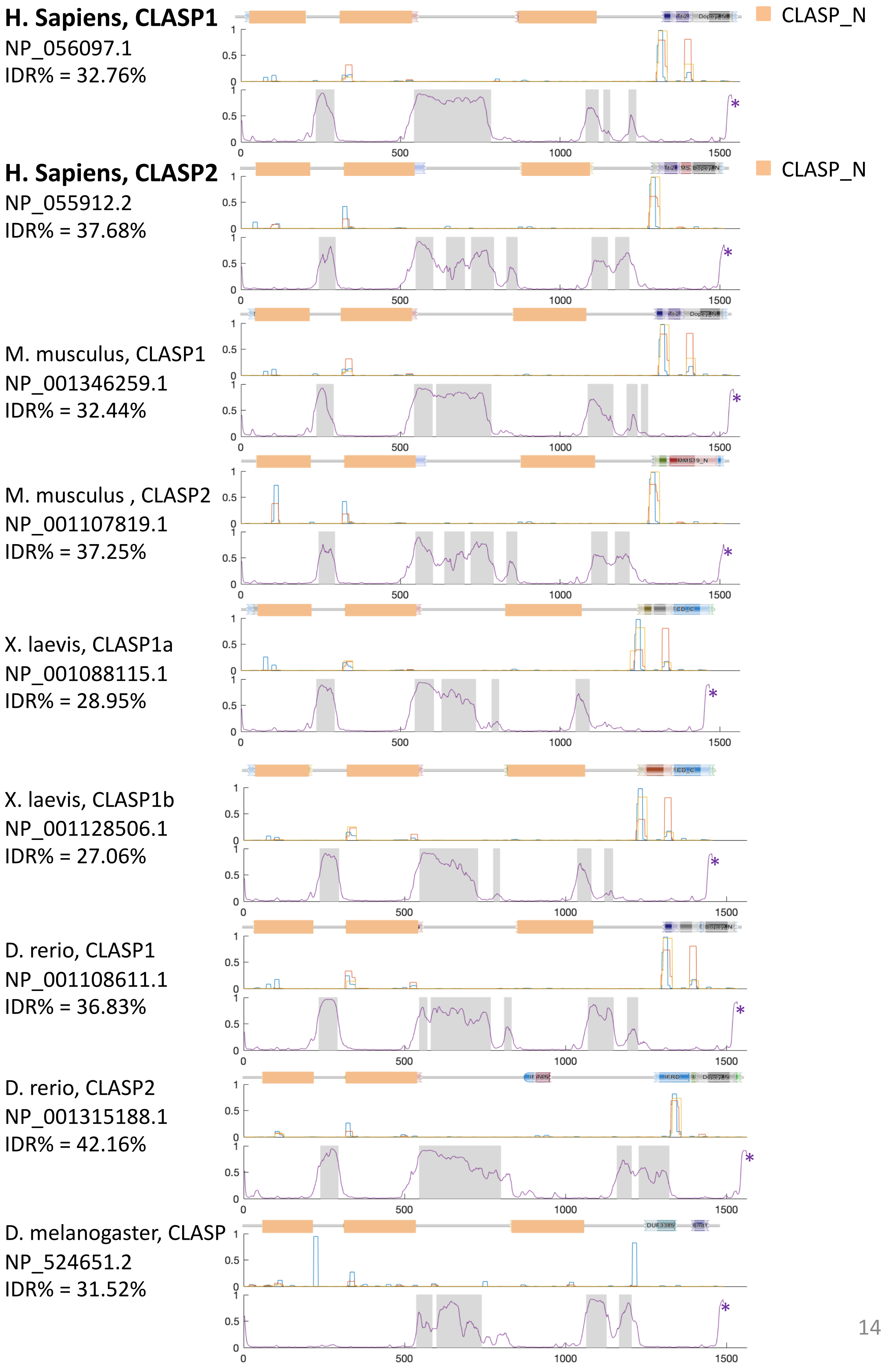
H. Sapiens, CLASP1
NP_056097.1
IDR% = 32.76%
CLASP_N
H. Sapiens, CLASP2
NP_055912.2
IDR% = 37.68%
CLASP_N
M. musculus, CLASP1
NP_001346259.1
IDR% = 32.44%
M. musculus , CLASP2
NP_001107819.1
IDR% = 37.25%
X. laevis, CLASP1a
NP_001088115.1
IDR% = 28.95%
X. laevis, CLASP1b
NP_001128506.1
IDR% = 27.06%
D. rerio, CLASP1
NP_001108611.1
IDR% = 36.83%
D. rerio, CLASP2
NP_001315188.1
IDR% = 42.16%
D. melanogaster, CLASP
NP_524651.2
IDR% = 31.52%

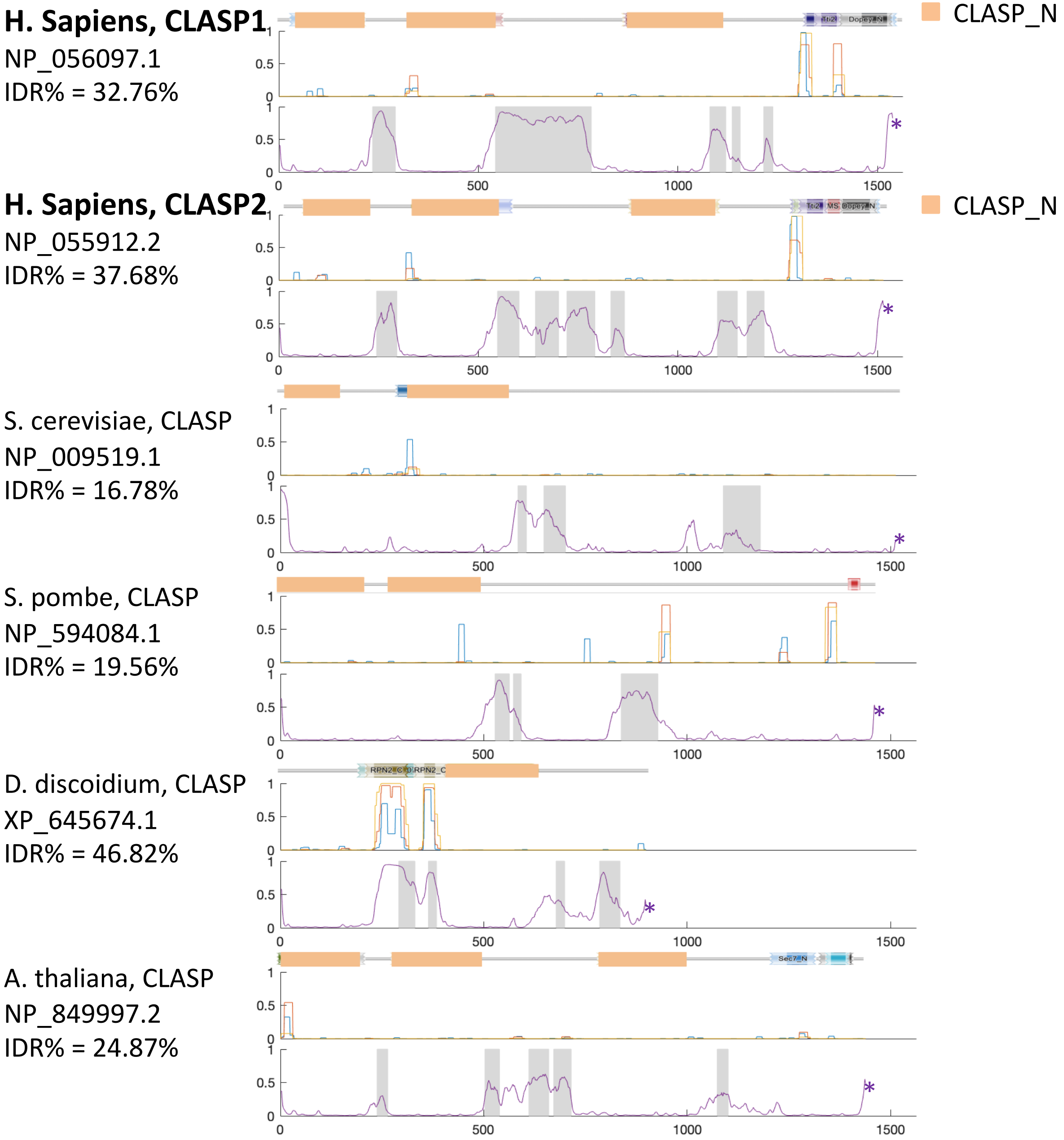
H. Sapiens, CLASP1
NP_056097.1
IDR% = 32.76%
CLASP_N
H. Sapiens, CLASP2
NP_055912.2
IDR% = 37.68%
CLASP_N
S. cerevisiae, CLASP
NP_009519.1
IDR% = 16.78%
S. pombe, CLASP
NP_594084.1
IDR% = 19.56%
D. discoidium, CLASP
XP_645674.1
IDR% = 46.82%
A. thaliana, CLASP
NP_849997.2
IDR% = 24.87%
0
500
1000
1500
0.5
1
*

**S8 Fig. Analysis of the position of coiled-coil domains and IDRs in CLASPs across a range of organisms.** For each CLASP protein (one chosen per species, accession number at left), the top image shows the domain structures identified by Pfam (e-value < 0.1). For ease of visualization, we manually colored the most significant domains (as indicated on the right), while minor domains are shown as plotted by Pfam. Note that a few recognized domain structures were not predicted with the e-value we selected, meaning that the absence of a domain in this figure should not necessarily be interpreted as absence of the domain in the protein. The middle image represents the probability of coiled-coil region as predicted by COILS with three different windows (7AA, blue; 14 AA, orange; 21 AA yellow). In the bottom image, the purple line shows the probability of IDRs predicted by Espritz, and the shaded area indicates the IDRs as predicted by MobiDB-lite. For both COILS and Espritz analyses, 1 indicates 100% likelihood. * indicates the end of the sequence.

### Reference:


1. Akhmanova A, Mausset-Bonnefont AL, van Cappellen W, Keijzer N, Hoogenraad CC, Stepanova T, et al. The microtubule plus-end-tracking protein CLIP-170 associates with the spermatid manchette and is essential for spermatogenesis. Genes Dev. 2005;19(20):2501-15.
2. van de Willige D, Hoogenraad CC, Akhmanova A. Microtubule plus-end tracking proteins in neuronal development. Cell Mol Life Sci. 2016;73(10):2053-77.
3. Hoogenraad CC, Koekkoek B, Akhmanova A, Krugers H, Dortland B, Miedema M, et al. Targeted mutation of Cyln2 in the Williams syndrome critical region links CLIP-115 haploinsufficiency to neurodevelopmental abnormalities in mice. Nat Genet. 2002;32(1):116-27.
4. Dix CI, Soundararajan HC, Dzhindzhev NS, Begum F, Suter B, Ohkura H, et al. Lissencephaly-1 promotes the recruitment of dynein and dynactin to transported mRNAs. J Cell Biol. 2013;202(3):479-94.
5. Geiser JR, Schott EJ, Kingsbury TJ, Cole NB, Totis LJ, Bhattacharyya G, et al. Saccharomyces cerevisiae genes required in the absence of the CIN8-encoded spindle motor act in functionally diverse mitotic pathways. Mol Biol Cell. 1997;8(6):1035-50.
6. Berlin V, Styles CA, Fink GR. BIK1, a protein required for microtubule function during mating and mitosis in Saccharomyces cerevisiae, colocalizes with tubulin. J Cell Biol. 1990;111(6 Pt 1):2573-86.
7. Giaever G, Chu AM, Ni L, Connelly C, Riles L, Veronneau S, et al. Functional profiling of the Saccharomyces cerevisiae genome. Nature. 2002;418(6896):387-91.
8. Yang C, Wu J, de Heus C, Grigoriev I, Liv N, Yao Y, et al. EB1 and EB3 regulate microtubule minus end organization and Golgi morphology. J Cell Biol. 2017;216(10):3179-98.
9. Elliott SL, Cullen CF, Wrobel N, Kernan MJ, Ohkura H. EB1 is essential during Drosophila development and plays a crucial role in the integrity of chordotonal mechanosensory organs. Mol Biol Cell. 2005;16(2):891-901.
10. Schwartz K, Richards K, Botstein D. BIM1 encodes a microtubule-binding protein in yeast. Mol Biol Cell. 1997;8(12):2677-91.
11. Bisgrove SR, Lee YR, Liu B, Peters NT, Kropf DL. The microtubule plus-end binding protein EB1 functions in root responses to touch and gravity signals in Arabidopsis. Plant Cell. 2008;20(2):396-410.
12. Pasqualone D, Huffaker TC. STU1, a suppressor of a beta-tubulin mutation, encodes a novel and essential component of the yeast mitotic spindle. J Cell Biol. 1994;127(6 Pt 2):1973-84.
13. Kirik V, Herrmann U, Parupalli C, Sedbrook JC, Ehrhardt DW, Hulskamp M. CLASP localizes in two discrete patterns on cortical microtubules and is required for cell morphogenesis and cell division in Arabidopsis. J Cell Sci. 2007;120(Pt 24):4416-25.
14. Wang PJ, Huffaker TC. Stu2p: A microtubule-binding protein that is an essential component of the yeast spindle pole body. J Cell Biol. 1997;139(5):1271-80.

15. Garcia MA, Vardy L, Koonrugsa N, Toda T. Fission yeast ch-TOG/XMAP215 homologue Alp14 connects mitotic spindles with the kinetochore and is a component of the Mad2-dependent spindle checkpoint. EMBO J. 2001;20(13):3389-401.
16. Guo L, Degenstein L, Dowling J, Yu QC, Wollmann R, Perman B, et al. Gene targeting of BPAG1: abnormalities in mechanical strength and cell migration in stratified epithelia and neurologic degeneration. Cell. 1995;81(2):233-43.
